# Manifold alignment for heterogeneous single-cell multi-omics data integration using Pamona

**DOI:** 10.1101/2020.11.03.366146

**Authors:** Kai Cao, Yiguang Hong, Lin Wan

## Abstract

Single-cell multi-omics sequencing data can provide a comprehensive molecular view of cells. However, effective approaches for the integrative analysis of such data are challenging. Although achieved state-of-the-art performance on single-cell multi-omics data integration and did not require any correspondence information, either among cells or among features, current manifold alignment based integrative methods are often limited by requiring that single-cell datasets be derived from the same underlying cellular structure. To overcome this limitation, we present Pamona, an algorithm that integrates heterogeneous single-cell multi-omics datasets with the aim of delineating and representing the shared and dataset-specific cellular structures. We formulate this task as a partial manifold alignment problem and develop a partial Gromov-Wasserstein optimal transport framework to solve it. Pamona identifies both shared and dataset-specific cells based on the computed probabilistic couplings of cells across datasets, and it aligns cellular modalities in a common low-dimensional space, while simultaneously preserving both shared and dataset-specific structures. Our framework can easily incorporate prior information, such as cell type annotations or cell-cell correspondence, to further improve alignment quality. Simulation studies and applications to four real data sets demonstrate that Pamona can accurately identify shared and dataset-specific cells, as well as faithfully recover and align cellular structures of heterogeneous single-cell modalities in the common space. Pamona software is available at https://github.com/caokai1073/Pamona.

## Introduction

The latest developments in high-throughput single-cell multi-omics sequencing technologies, e.g., single-cell RNA-sequencing (scRNA-seq) and ATAC-sequencing (scATAC-seq), enable cell-resolved investigation of heterogeneous cellular populations that make up tissues, the dynamics of developmental processes, and the underlying regulatory mechanisms that control cellular functions [1]. The integration of emerging single-cell multi-omics datasets, however, poses fresh data integration challenges such as unmatched/distinct features and/or unpaired cells across datasets [2]. Integrative methods have been developed to enable joint learning across multiple types of data. For example, the celebrated single-cell data analysis platform Seurat [3] projected (distinct) feature spaces across datasets into a common subspace using canonical correlation analysis (CCA), which maximizes inter-dataset correlation, and selected mutual nearest-neighbors (MNNs) [4] as anchors to align datasets. Although it achieved success in batch effect correction, Seurat relies on the linear mapping of CCA and the linear alignments of MNNs, thus weakening its ability to handle nonlinear geometrical deformations and rotations of intrinsic manifolds embedded across cellular modalities. MOFA+ employed a Bayesian Group Factor Analysis approach, and scAI [5] adopted a non-negative matrix factorization (NMF) approach for integrating single-cell multi-omics datasets with distinct features. However, both required datasets of paired cells.

Recently, manifold alignment approaches, which aimed to align embedded low-dimensional manifolds, have been developed for holistic representation of the intrinsic cellular structures across cellular modalities, without requiring any correspondence information, either among cells or among features, e.g., MATCHER [6], MMD-MA [7, 8], UnionCom [9], and SCOT [10]. These methods were derived under various advanced machine learning techniques, such as linear trajectory alignment using the latent Gaussian process, as in MATCHER [6]; kernel space matching based on maximum mean discrepancy, as in MMD-MA [7]; metric space matching based on the graph-matching/quadratic assignment formulation, as in UnionCom [9], or the optimal transport formulation, as in SCOT [10]. Although these state-of-the-art methods have achieved integrative performance with encouraging results [10], current manifold alignment methods often automatically assume that all datasets share the same underlying structure across cellular modalities. Such assumption can be easily nullified by presenting dataset-specific cell types/structures across the single-cell datasets. Therefore, it remains computationally challenging for state-of-the-art manifold alignment algorithms to preserve both shared and dataset-specific cellular structures across datasets during integration.

Here, we present Pamona, a <u>pa</u>rtial Gro<u>mo</u>v-Wasserstei<u>n</u> based manifold <u>a</u>lignment algorithm, that integrates heterogeneous single-cell multi-omics datasets to delineate and represent both shared and dataset-specific cellular structures (Fig. 1a and Methods). Gromov-Wasserstein (GW) distance, a generalized optimal transport which overcomes the lack of intrinsic correspondence between feature spaces, has been proven to be increasingly valuable for diverse fields [11, 12]. In the single-cell data analysis community, GW has been applied to the spatial reconstruction of gene expression cartography by novoSpaRc [13] and single-cell multi-omics data integration by SCOT [10]. While GW seeks a transportation map that preserves the total mass between the two probability distributions [11, 12], partial-GW extends the GW framework by allowing only a fraction of the total mass to be transported [14]. The spirit of partial-GW is built upon adding virtual or dummy points onto the marginals and enforcing points with large discrepancies absorbed by the virtual points [14, 15]. As such, partial-GW enables Pamona to reconstruct the probabilistic couplings of cells across datasets to identify both shared and dataset-specific cells. Based on the probabilistic couplings, Pamona further aligns single-cell multi-omics datasets in a common low-dimensional space, while preserving both shared and dataset-specific cellular structures across modalities.

**Figure 1:**
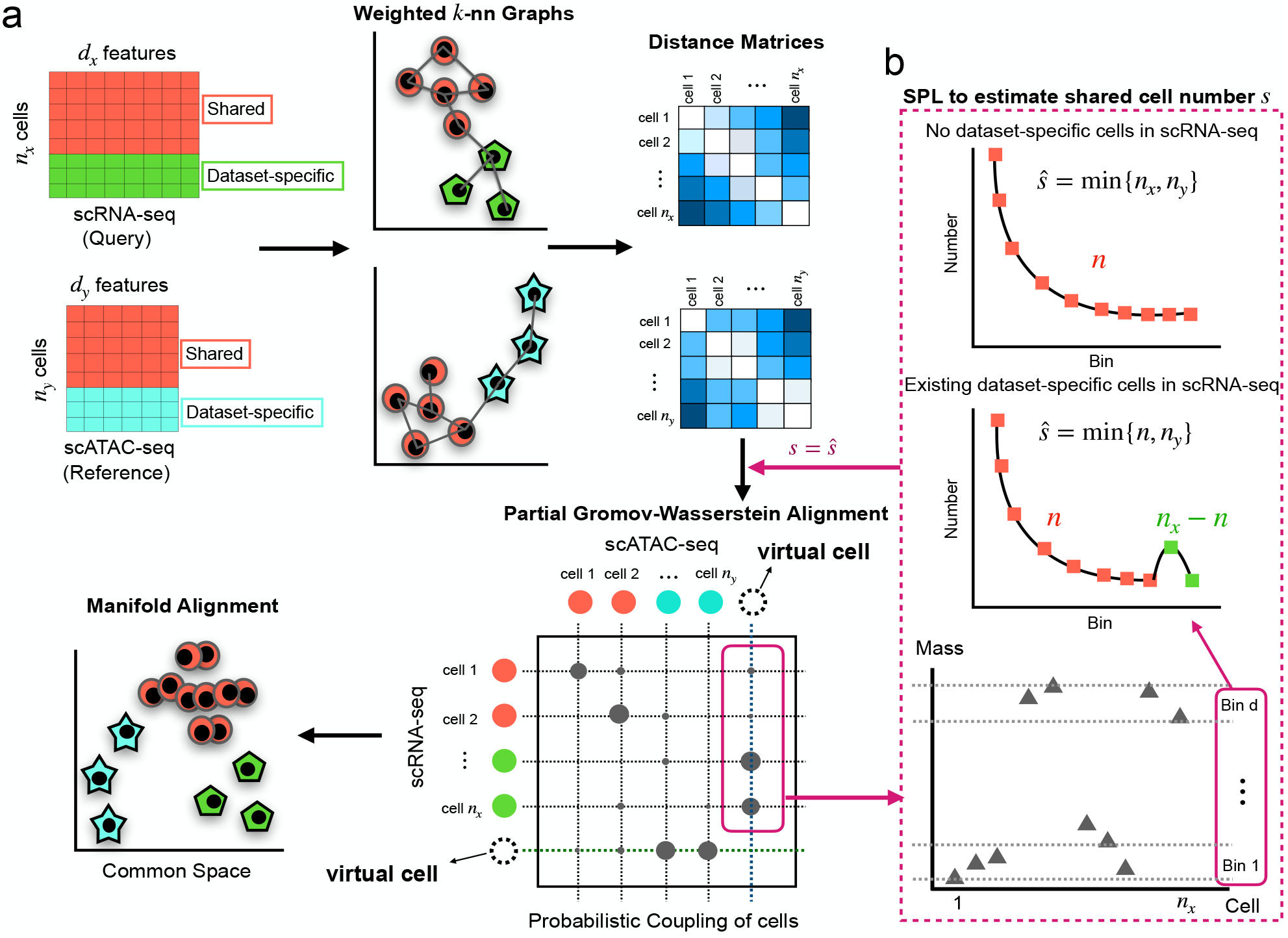
Overview of Pamona. Pamona is a partial manifold alignment algorithm for heterogeneous single-cell multi-omics data integration. Given inputs of multiple cellular modalities (e.g., scRNA-seq and scATAC-seq), it identifies both shared and dataset-specific cells based on the computed probabilistic couplings of cells across datasets, and it aligns cellular modalities in a common low-dimensional space, while simultaneously preserving both shared and dataset-specific structures. (a) Pamona constructs a weighted *k*-nn graph of cells for each dataset (step 1), computes the geodesic distance matrix of cells within each dataset (step 2), computes the probabilistic coupling matrices of cells based on the partial Gromov-Wasserstein optimal transport (step 3), and aligns cellular modalities with distinct unmatched features in a common low-dimensional space to holistically represent the cellular structures (step 4). (b) A Scree-Plot-Like (SPL) method is proposed to estimate the shared cell number *s* when it is not available.

Before Pamona, only a few methods were developed specifically for integrating single-cell datasets with dataset-specific cell types/structures. For example, Scanorama efficiently integrated multiple scRNA-seq datasets based on a generalized MNN matching technique for “panorama stitching” of heterogeneous scRNA-seq datasets. Liger [16], which employed an integrative NMF approach to find the shared and dataset-specific components across datasets, required pre-matched common feature space across modalities and could not integrate datasets into a common space. The manifold alignment method UnionCom [9] showed its ability to accommodate dataset-specific cells, but remains to be further explored.

Notably, Pamona can perform both global and partial manifold alignments for single-cell multiomics data integration. In this study, we propose a Scree-Plot-Like (SPL) method paralleled with Pamona to estimate the shared cell number which needs to be specified by the partial-GW framework (Fig. 1b). With no inherent reliance on any prior information, our framework offers the flexibility to match prior information, e.g., cell type annotations or cell-cell correspondence, when available. To assess its performance, we applied Pamona to 2 simulated and 4 real single-cell multiomics data sets on various tasks. We demonstrate that Pamona can accurately identify shared and dataset-specific cells, as well as faithfully recover and align intrinsic manifolds across heterogeneous cellular modalities in the common space.

## Results

### Overview of Pamona

Pamona is a partial manifold alignment algorithm for heterogeneous single-cell multi-omics data integration. The main inputs of Pamona are the data matrices of single-cell multimodal profiles, e.g., gene expression, chromatin accessibility, and DNA methylation. The main outputs of Pamona are (1) the probabilistic couplings of cells across datasets in order to identify both shared and dataset-specific cells and (2) the common low-dimensional space that recovers and aligns intrinsic structures of heterogeneous cellular modalities.

As illustrated in Fig. 1a, Pamona integrates single-cell multi-omics datasets in 4 major steps as follows. (1) It constructs a weighted *k* nearest neighbor (*k*-nn) graph of cells for each dataset based on Euclidean distance. (2) It computes the geodesic distance matrix of cells within each dataset based on the pairwise shortest distances between nodes/cells on the *k*-nn graph. (3) It computes the probabilistic coupling matrices of cells based on the partial-GW optimal transport to identify cell-cell correspondence across cellular modalities. (4) It aligns cellular modalities with distinct unmatched features in a common low-dimensional space for feature comparability and the holistic representation of cellular structures.

The Pamona analytical framework is mainly derived from partial-GW optimal transport [14] (Methods). The partial-GW framework needs to specify the shared cell number *s* which is often unavailable in practice. Therefore, we propose the SPL method to accurately estimate the shared cell number (Fig. 1b and Methods). Besides, Pamona does not rely on prior information like cell types or cell-cell correspondence, but it does offer the flexibility to match prior knowledge when available (Methods).

### Pamona improved heterogeneous single-cell multi-omics data integration

In the following, we compared Pamona to prior state-of-the-art single-cell multi-omics integration methods, including Seurat v3 [3], MMD-MA [7], UnionCom [9], and SCOT [10].

For this purpose, we employed 2 simulated and 4 real-world single-cell multi-omics data sets as follows: a simulated data set from SCOT [10], hereinafter denoted as Simulation 1; a simulated data set from UnionCom [9], hereinafter denoted as Simulation 2; the single-cell analysis of genotype, expression and methylation data set from [17], hereinafter denoted as sc-GEM; the single-cell nucleosome, methylome and transcriptome data set from [18], hereinafter denoted as scNMT-seq; the single-nucleus chromatin accessibility and mRNA expression data set from [19], hereinafter denoted as SNARE-seq; and the 10X Genomics scRNA-seq and scATAC-seq data set of human peripheral blood mononuclear cells from [20], hereinafter denoted as PBMC. Detailed information of these data sets is provided in Methods.

We mainly employed two scores to assess the performance of single-cell multi-omics data integration: (i) Label Transfer Accuracy to measure the ability to transfer labels of the shared cells from one dataset to another and (ii) Alignment Score to measure the ability to preserve both shared and dataset-specific structures. In addition, we adopted the FOSCTTM score to measure the preservation of cell-cell correspondence across datasets for the SNARE-seq data set. All three scores work on the basis of the common space by integrative methods (See Methods for details).

In general, Pamona improved various partial manifold alignment tasks with the highest scores of Label Transfer Accuracy and Alignment Score on Simulation 1, Simulation 2, sc-GEM, and scNMT-seq (Supplementary Fig. 1). Especially, Pamona increased the Alignment Score markedly on the partial manifold alignment tasks. On the global manifold alignment task of the scNMT-seq data set, Pamona achieved the second highest performance, slightly below the highest one achieved by SCOT (Supplementary Fig. 1d). Detailed results and comparison will be provided in the following sections.

Meanwhile, Pamona is robust to the hyperparameter choices: the number of neighborhoods *k* of the *k*-nn graph, the regularization parameter *E* of the partial-GW framework, the parameter *λ* of manifold alignment to make a trade-off between aligning corresponding cells and preserving the local geometries, and the cost *α* of the virtual points (Supplementary Fig. 2).

### Pamona resolved the partial manifold alignment on simulated data sets

We assessed the performance of Pamona on partial manifold alignment using Simulation 1 and Simulation 2.

In Simulation 1, dataset **X** contains an embedded bifurcated tree with three branches denoted as Type 1 (blue points), Type 2 (green points), and Type 3 (red points), respectively (Supplementary Fig. 3a). The original dataset **Y** also contains an embedded bifurcated tree with cell types corresponding to those in Dataset **X**. To construct a partial manifold alignment task, we removed the cells of Type 3 from dataset **Y**, resulting in a lineage structure constituted only by cells of Type 1 (blue points) and Type 2 (green points) (Supplementary Fig. 3a). Therefore, Type 3 becomes an **X**-specific branch. When we applied Pamona to integrate **X** and **Y**, it aligned the shared cells of Type 1 (blue points) and Type 2 (green points), accordingly, and preserved the cells of Type 3 as the **X**-specific branch in the common space (Supplementary Fig. 3b), achieving the highest Alignment Score of 0.834 and Label Transfer Accuracy of 97.92%. In contrast, Seurat v3, MMD-MA, UnionCom, and SCOT did not preserve either shared or dataset-specific structures very well (Supplementary Fig. 4), resulting in sharp drops of Alignment Score (Seurat v3, 0.215; MMD-MA, 0.507; UnionCom, 0.596; SCOT, 0.601) (Supplementary Fig. 1a). Meanwhile, MMD-MA, SCOT and UnionCom achieved Label Transfer Accuracy similar to that of Pamona (MMD-MA, 95.85%; SCOT, 95.02%; UnionCom, 97.91%), while Seurat v3 had the lowest Label Transfer Accuracy of 70.12% (Supplementary Fig. 1a).

In Simulation 2, dataset **X** contains an embedded bifurcated tree with three branches denoted as Type 1 (red points), Type 2 (blue points), and Type 3 (yellow points), respectively; dataset **Y** contains an embedded trifurcated tree with three branches corresponding to those in dataset **X** and a **Y**-specific branch of cells from Type 4 (green points) (Supplementary Fig. 3c). When we applied Pamona to integrate **X** and **Y**, it aligned the shared cells of Type 1 (red points), Type 2 (blue points) and Type 3 (yellow points), accordingly, and preserved the cells of Type 4 as the **Y**-specific branch in the common space (Supplementary Fig. 3d). It had the highest Alignment Score of 0.885 and Label Transfer Accuracy of 93.5% (Supplementary Fig. 1b). In comparison, UnionCom also integrated **X** and **Y** quite well (Supplementary Fig. 5b) and had the second highest Alignment Score of 0.702 and Label Transfer Accuracy of 90.5% (Supplementary Fig. 1b). In contrast, Seurat v3, MMD-MA, and SCOT did not perform well on the partial manifold task (Supplementary Fig. 5), showing sharp drops of Alignment Score (Supplementary Fig. 1b).

### Pamona identified informative genes in delineating the shared and dataset-specific cellular structures of the sc-GEM data set

We applied Pamona to the sc-GEM data set of gene expression and DNA methylation on samples of human cells undergoing reprogramming to induced pluripotent stem (iPS) cells (Methods). In a previous study, we applied UnionCom to this same data set for a global manifold alignment task [9]. However, in this study, we removed the human foreskin fibroblast (BJ) cells from the DNA methylation dataset to construct a partial manifold alignment task. Both gene expression and DNA methylation datasets demonstrated similar linear structures with the same cell type orders when visualized using Uniform Manifold Approximation and Projection (UMAP) [21, 22] separately (Fig. 2a). The gene expression dataset showed the dataset-specific cell type BJ (green points) located at one end of the linear trajectory (Fig. 2a, lower panel).

**Figure 2:**
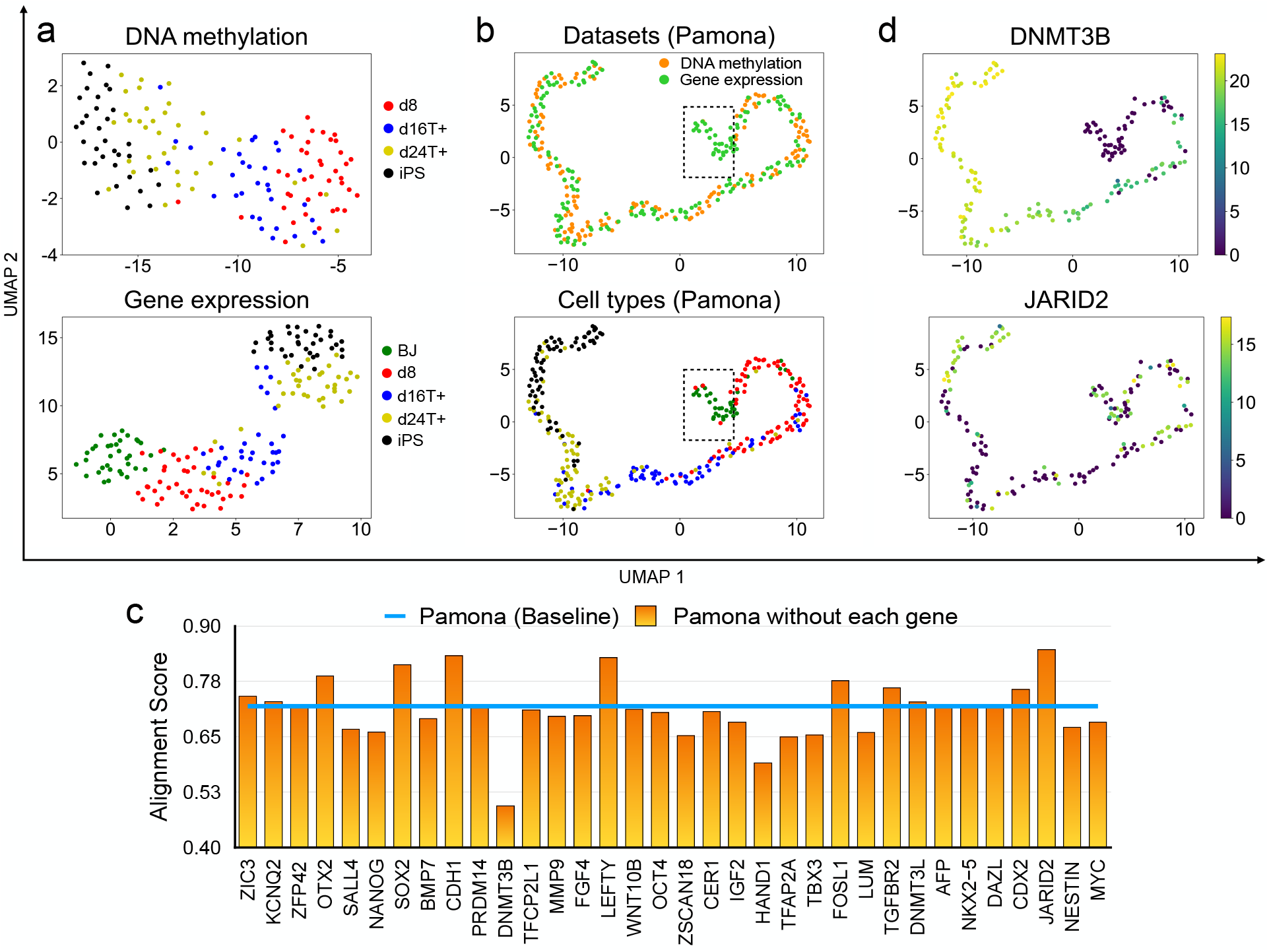
Pamona integrated the sc-GEM data set and identified informative genes in delineating the shared and dataset-specific cellular structures. (a) Visualizations of the DNA methylation (upper panel) and gene expression (lower panel) datasets separately using UMAP before alignment. BJ cells (green points) comprise the gene expression dataset-specific cells. (b) Visualizations of the common space of the two aligned datasets by Pamona using UMAP: upper panel: cells are colored according to their corresponding datasets; lower panel: cells are colored according to their corresponding types. (c) Alignment score of Pamona using expression profile with all 34 genes (blue line, baseline) and without each of the 34 genes separately (orange bars). (d) Gene expression of cells in the common space: upper panel: DNMT3B (informative gene); lower panel: JARID2 (non-informative gene).

Pamona aligned the shared cells of d8 (red points), d16T+ (blue points), d24T+ (yellow points) and iPS (black points), accordingly, and preserved BJ cells as the gene expression dataset-specific cells in the common space (Fig. 2b). It achieved the highest Alignment Score of 0.719 and Label Transfer Accuracy of 66.2% (Supplementary Fig. 1c). In comparison, UnionCom achieved the second highest Alignment Score of 0.592 and Label Transfer Accuracy of 45.77% (Supplementary Fig. 1c). SCOT and UnionCom did not separate BJ cells from d8 cells (Supplementary Fig. 6a,b); MMD-MA and Seurat v3 failed to align the shared cells across the two datasets (Supplementary Fig. 6c,d) and showed relatively low accuracy (Supplementary Fig. 1c).

We futher assessed the importance of all 34 genes from the gene expression profile in delineating the shared and dataset-specific cellular structures. We set the Alignment Score achieved by Pamona at 0.719 as baseline (Fig. 2c, blue line). We removed each gene from the gene expression profile, applied Pamona, and computed the Alignment Score separately (Fig. 2c, orange bars). We found that (1) removing each of DNMT3B, HAND1, TFAP2A and TBX3 genes resulted in a significant lose of Alignment Score, suggesting that these genes are informative features for the partial alignment task, but that (2) removing each of the CDH1, LEFTY and JARID2 genes resulted in a significant gain of Alignment Score, suggesting that these genes are non-informative features for the partial alignment task. It can be seen in the figure that the informative gene DNMT3B was highly expressed in the shared cells, but not expressed in BJ cells, which are dataset-specific cells (Fig. 2d, upper panel). The non-informative gene JARID2 was uniformly expressed in both shared and dataset-specific cells (Fig. 2d, lower panel).

### Pamona resolved the integration of the scNMT-seq data set in both global and partial manifold alignment tasks

We applied Pamona to the scNMT-seq data set of chromatin accessibility, DNA methylation and gene expression on mouse gastrulation samples collected at four time stages, i.e., embryonic day 4.5 (E4.5), E5.5, E6.5 and E7.5 [18].

We conducted a global manifold alignment task to integrate the two cellular modalities of chromatin accessibility and DNA methylation. Both datasets demonstrated similar linear structures which preserve the time stage orders when visualized using UMAP separately (Fig. 3a). Pamona aligned the two datasets in a common space and also preserved the time stage orders (Fig. 3b). Pamona achieved the second highest Alignment Score of 0.866 and Label Transfer Accuracy of 70.79%, slightly below the highest scores achieved by SCOT (Alignment Score 0.88; Label Transfer Accuracy 74.03%). See Supplementary Fig. 7a-d and Supplementary Fig. 1d for more results of the 4 compared methods.

**Figure 3:**
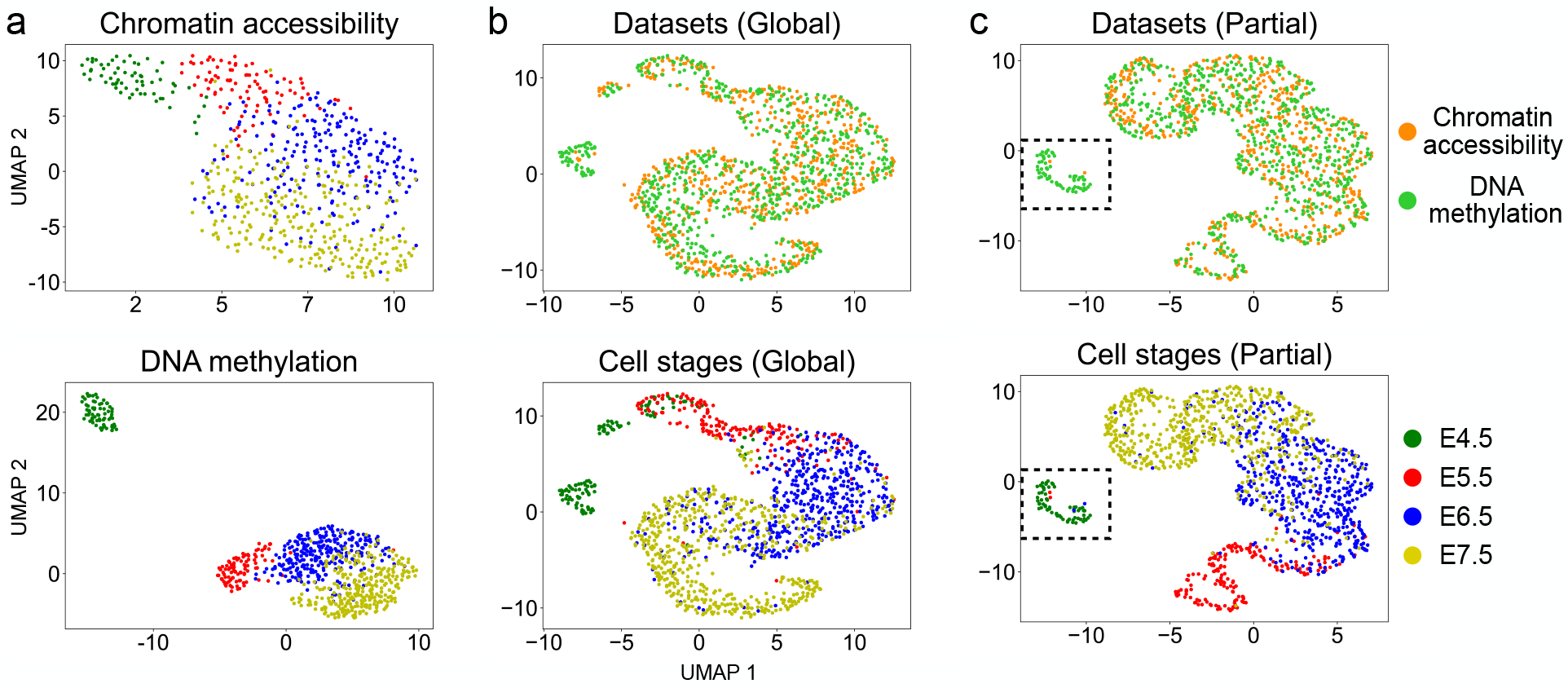
Pamona integrated the scNMT-seq data set in both global and partial manifold alignment tasks. (a) Visualizations of chromatin accessibility (upper panel) and DNA methylation (lower panel) datasets separately using UMAP before alignment. (b) Visualizations of the common space of the global alignment of the two datasets by Pamona using UMAP: upper panel: cells are colored according to their corresponding datasets; lower panel: cells are colored according to their corresponding types. (c) Visualizations of the common space of the partial alignment of the two datasets (cells of E4.5 were removed from the chromatin accessibility dataset) by Pamona using UMAP.

Next, we constructed a partial manifold alignment task by removing the cells of time stage E4.5 from the chromatin accessibility dataset. Pamona aligned the shared cells from E5.5 to E7.5, accordingly, and preserved the cells of E4.5 as the DNA methylation dataset-specific cells in the common space (Fig. 3c). It achieved the highest Alignment Score of 0.907 and Label Transfer Accuracy of 75.34% (Supplementary Fig. 1e). In comparison, UnionCom achieved the second highest Alignment Score of 0.669 and Label Transfer Accuracy of 64.72%. SCOT, which was designed with the underlying assumption of global manifold alignment, dropped its accuracy markedly (Alignment Score 0.297; Label Transfer Accuracy 62.67%). See Supplementary Fig. 7e-h and Supplementary Fig. 1e for more results of the 4 compared methods.

Finally, we conducted a partial manifold alignment task to integrate the three cellular modalities of chromatin accessibility, gene expression and DNA methylation by removing the cells of time stage E4.5 from the chromatin accessibility dataset (Supplementary Fig. 8a). Pamona successfully integrated the three modalities by aligning the shared cells according to their time stages and preserving the cells of E4.5 as the DNA methylationand gene expression-specific cells in the common space (Supplementary Fig. 8b). In contrast, UnionCom, which can also handle multiple datasets, did not clearly separate the cells of E4.5 of the DNA methylation and gene expression datasets from the cells of the chromatin accessibility dataset (Supplementary Fig. 8c).

### Pamona improved single-cell multi-omics data integration by incorporating partial cell-cell correspondence information

We assessed the performance of Pamona on integration of the SNARE-seq data set by incorporating partial cell-cell correspondence information. The SNARE-seq data set of gene expression and chromatin accessibility was derived from samples of mixed cultured human BJ, H1, K562 and GM12878 cells [19], and it was previously tested with SCOT [10]. Since the gene expression and chromatin accessibility profile was sequenced from the same cells, cell-cell correspondence information was available for the SNARE-seq data set.

We followed the procedure of SCOT [10] to visualize the structures of the two cellular modalities of gene expression and chromatin accessibility using principal component analysis (PCA) separately (Supplementary Fig. 9a). When no prior information was utilized, SCOT slightly outperformed Pamona in preserving the accuracy of cell-cell correspondence, as indicated by the FOSCTTM scores of 0.156 (SCOT) and 0.157 (Pamona) (Supplementary Fig. 9b).

However, when incorporating partial cell-cell correspondence information, Pamona showed improvement in both preserving the alignment accuracy of cell-cell correspondence (Supplementary Fig. 9c) and maximizing computational efficiency (Supplementary Fig. 9d). For example, when incorporating cell-cell correspondence of 30%, 60% and 90%, the FOSCTTM scores by Pamona dropped markedly to 0.15, 0.132, and 0.116, respectively (Supplementary Fig. 9c). By incorporating partial cell-cell correspondence, Pamona showed up to 10-fold acceleration of computational speed, e.g., Pamona with 30% cell-cell correspondence is about five times faster than Pamona without the information (Supplementary Fig. 9d).

### Pamona resolved the integration of the heterogeneous PBMC data set by incorporating cell type annotation information

We applied Pamona to integrate the heterogeneous PBMC data set by incorporating cell type annotation information. The PBMC data set consisting of gene expression (scRNA-seq) and chromatin accessibility (scATAC-seq) was derived from samples of human peripheral blood mononuclear cells released by 10X Genomics, and it was previously analyzed by MAESTRO [20]. We adopted the annotations provided by MAESTRO [20] as the benchmark to assess the performance of the methods (see Fig. 4a for the annotated cell types of scRNA-seq and scATAC-seq datasets on UMAP visualizations separately). Since both scRNA-seq and scATAC-seq have dataset-specific cell types annotated by MAESTRO (Fig. 4a), such data integrative analysis is a partial manifold alignment task.

**Figure 4:**
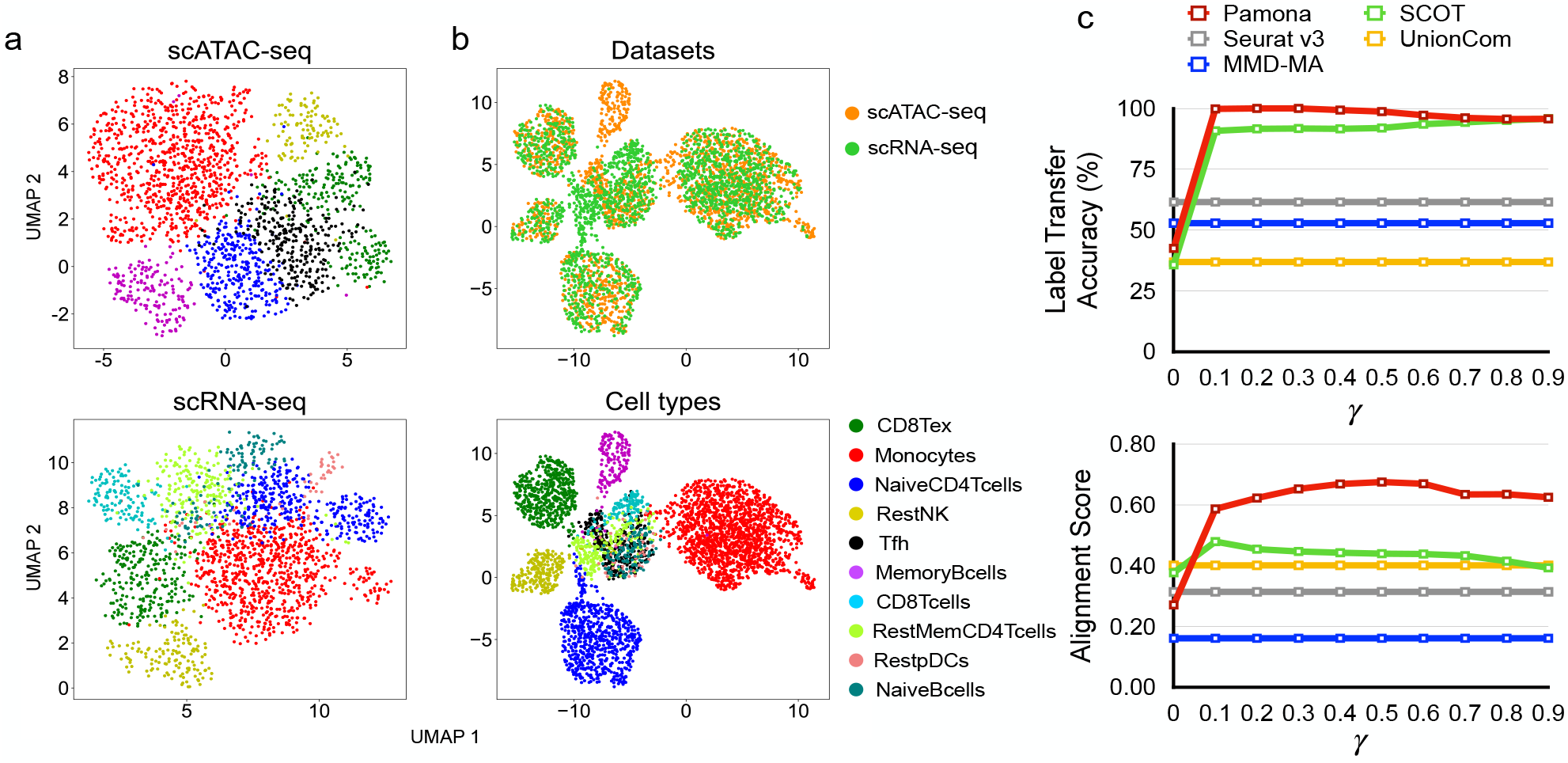
Pamona integrated the heterogeneous PBMC data set by incorporating cell type annotation information. (a) Visualizations of the scATAC-seq (upper panel) and scRNA-seq (lower panel) datasets separately using UMAP before alignment. (b) Visualizations of the common space of the two aligned datasets by Pamona by incorporating cell type annotation information (*γ* = 0.5): upper panel: cells are colored according to their corresponding datasets; lower panel: cells are colored according to their corresponding types. (c) The Label Transfer Accuracy and Alignment Score of Pamona and SCOT when *γ* in the disagreement matrix **M** was set from 0 to 0.9, which is equivalent to the growing influence of cell annotations compared to data structure. We similarly incorporated prior information for SCOT. Results by Seurat v3, MMD-MA and UnionCom did not incorporate cell type annotation information.

When no prior information was used, all five methods tested had low scores of Label Transfer Accuracy and Alignment Score (Fig. 4c, when *γ* = 0). None of the 5 methods tested could integrate the two modalities since they failed to preserve both shared and dataset-specific cellular structures (Supplementary Fig. 10). For example, Pamona could not align shared NaiveCD4T cells across the two datasets in the common space (Supplementary Fig. 10).

When incorporating cell type annotation information, however, Pamona aligned the shared cells, accordingly, and preserved the dataset-specific cells in the common space (Fig. 4b) with demonstrably improved integrative accuracy in that the scores of Label Transfer Accuracy and Alignment Score increased as the *γ* increased (Fig. 4c). We also incorporated cell type annotation information for SCOT, as we did for Pamona, but SCOT only slightly improved its Alignment Score (Fig. 4c) since it lacks a partial manifold alignment strategy to separate out dataset-specific cells from the shared cells in the common space.

## Discussion

In this study, we propose Pamona, a partial manifold alignment algorithm, for heterogeneous single-cell multi-omics data integration. Pamona delineates and represents the shared and dataset-specific cell structures in the common space across modalities. It easily incorporates prior information, such as cell type annotations or cell-cell correspondence, to further improve alignment quality. When applied to 2 simulated and 4 real single-cell multi-omics data sets, Pamona accurately identified shared and dataset-specific cells, and it faithfully recovered and aligned cellular structures of heterogeneous cellular modalities in the common space.

Pamona was developed based on the recently proposed partial-GW optimal transport framework [14]. The key technique of partial-GW is adding virtual/dummy points onto the marginals to enforce points with large discrepancies absorbed by the virtual points [14, 15]. Virtual points have also been discussed in the partial graph matching problem [23]. As we can see from the computed probabilistic coupling matrices by Pamona, the virtual points achieved the goal of absorbing the dataset-specific cells (Supplementary Figs. 11-13). We also proposed an SPL method to estimate the shared cell number across datasets. We demonstrate that SPL is very accurate in our tested datasets (Supplementary Fig. 14).

Pamona is a computationally efficient algorithm. However, it requires *O*(*n*^2^) memory consumption in the storage of distance matrices. Therefore, it may not perform well when sample size is in large-scale (e.g., *>* 1 million cells). As large-scale single-cell multi-omics datasets are emerging, it is challenging to resolve the scalability problem for Pamona. One approach to tackle this problem is to develop a distributed storage and distributed computational framework for Pamona. Meanwhile, since large-scale single-cell datasets can be highly redundant, we can take the alternative approach by (1) adopting the state-of-the-art neural network with mini-batch framework [24], or (2) selecting a subset of informative samples using the advanced geometric sketching tool [25] prior to applying Pamona. We plan to pursue these topics in our future work.

## Methods

### Mathematical formulation of Pamona

Pamona is a partial manifold alignment algorithm for single-cell multi-omics data integration. The procedure used by Pamona includes 4 major steps (see Fig. 1a and Supplementary Note 1 for the pseudocode).

First, suppose that 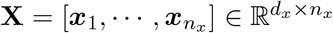 and 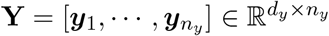 are the inputs of two single-cell multi-omics datasets where *d*_*x*_(*d*_*y*_) and *n*_*x*_(*n*_*y*_) are the number of features and cells for **X**(**Y**). We compute the weighted *k*-nn graphs for each of the two datasets where the nodes of each graph correspond to cells within the dataset, and edges have weights based on pairwise Euclidean distances between cells. In case the *k*-nn graph for a given *k* is not connected, we adopt the same procedure as that in Klimovskaia et al. [26] to enforce connectivity.

Second, we compute the geodesic distances of cells within the same dataset by calculating the shortest distance between each pair of nodes (cells) on the *k*-nn graph using the Dijkstra algorithm. The path with the shortest distance will approximate to geodesic distance on the embedded manifold [27]. We denote the geodesic distance matrices for **X** and **Y** as 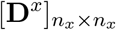 and 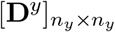, respectively.

Third, we compute the probabilistic cell-cell correspondence between **X** and **Y** to identify the shared and dataset-specific cells. Here, we formulate the problem as the partial-GW optimal transport framework [14]. Partial-GW extends the GW optimal transport to allow only a fraction of the total mass to be matched/transported [12].

Specifically, we assign each cell from each of the two datasets with a point mass 1*/N*, where *N* = max{*n*_*x*_, *n*_*y*_}. Partial-GW aims to match (transport) a fraction of *s/N* mass from **X** to **Y**. Here, *s* ≤ min{*n*_*x*_, *n*_*y*_} needs to be specified, and it can be regarded as the number of shared cells between **X** and **Y**. Partial-GW finds a probabilistic coupling matrix 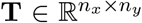 from *n*_*x*_ cells in **X** to *n*_*y*_ cells in **Y** able to minimize discrepancy between the geodesic distances in 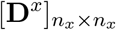 and 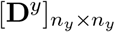, that is

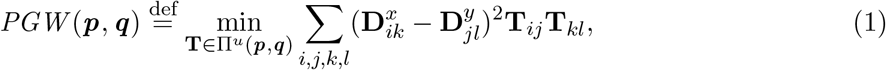

where **T**_*ij*_ is the relative probability that matches cell *i* in **X** to cell *j* in **Y**, satisfying the constraints on the set of all admissible coupling Π^*u*^(***p, q***) as

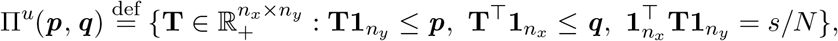

where 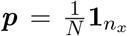 and 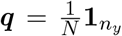 are the mass marginal distributions for **X** and **Y**, respectively. Here, **1**_*n*_ ∈ ℝ ^*n*^ denotes an *n*-dimensional vector of ones, and the superscript T denotes the transpose of a vector or matrix. The equality 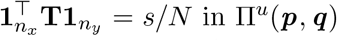 in Π^*u*^ (***p, q***) enforces the relaxed requirement that only a fraction of *s/N* cells needs to be matched/transported between the two datasets.

We write *PGW* in matrix form and add an entropic regularization penalty to the original problem, resulting in the entropic regularized partial-GW metric as follows:

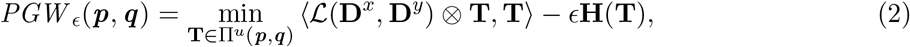

where ⟨·, ·⟩ denotes Frobenius dot product of matrices, (*ℒ* ⊗ **T**) denotes an *n*_*x*_ × *n*_*y*_ cost matrix with its (*i, j*)-th element defined as 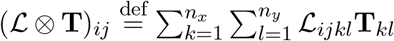, the discrepancy between geodesic distances 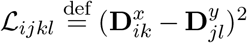, the entropic regularization term 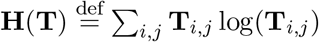, and *ϵ* is a tradeoff parameter between *PGW* and **H**(**T**).

To solve the optimization problem of *PGW* _*ϵ*_, Pamona adds virtual points onto the marginals as in [14]. The virtual points are used as buffers when comparing distributions with different probability masses. In this way, the partial-GW problem is equivalent to a point (cell) augmented, but still standard GW problem, which can be efficiently solved by Sinkhorn iterations [28, 29]. Pamona solves it iteratively as follows: for each iteration *k*:

A1. Update the cost matrix **C**^(*k*)^ = (*ℒ* ⊗ **T**)^(*k*)^ as follows:

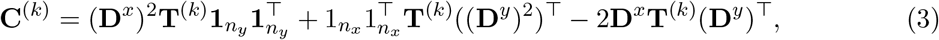

where 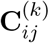 represents the cost of aligning cell *i* in **X** to cell *j* in **Y** at iteration *k*.

A2. Add two virtual points (cells), one to **X** and the other to **Y**, resulting in augmented cost matrix 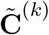and marginal distributions 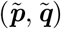 defined as follows:

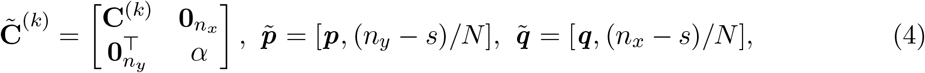

where the variable *α* ∈ *R*_+_ is set as a relatively large value greater than the elements of the cost matrix **C**^(*k*)^, with the aim of preventing the alignment within virtual cells between two datasets. In practice, *α* can be chosen as any value such that 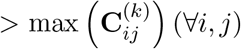, and the performance of Pamona is robust to the choice of *α* (see Supplementary Fig. 2e,f). The mass of virtual cell in **X** is set as (*n*_*y*_ − *s*)*/N* in 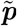, and the mass of virtual cell in **Y** is set as (*n*_*x*_ − *s*)*/N* in 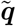.

A3. Compute GW optimal transport plan with 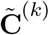 and 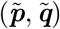. We first normalize 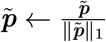 and 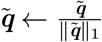 to construct probability distributions. Afterwards, we formulate the problem as

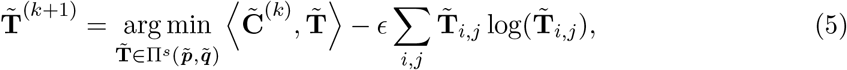

where

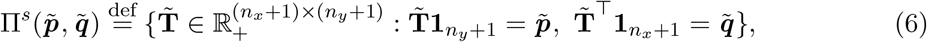

which is a standard entropic regularized optimal transport problem. The 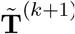 is efficiently solved by Sinkhorn iterations [29]. Once 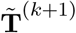 is obtained, we remove the last row and column of 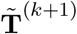 to obtain **T**^(*k*+1)^ as in [14].

The mechanism of adding virtual points with the designed augmented cost matrix 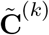 and marginal distributions 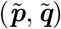, as defined above, is based on the theory that the virtual cell in **X** attracts mass of (*n*_*y*_ − *s*)*/N* cells in **Y**, with large values in corresponding columns of the cost matrix 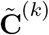, and that the virtual cell in **Y** attracts mass of (*n*_*x*_ − *s*)*/N* cells in **X**, with large values in corresponding rows of the cost matrix 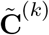.

Four, we then perform manifold alignment by aligning cellular modalities with distinct un-matched features in a common low-dimensional space for feature comparability. The common space should preserve both shared and dataset-specific structures across cellular modalities. Suppose we have *l* + 1(*l* ≥ 1) datasets. As in Seurat [3], we fix a dataset 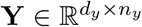as the reference dataset, and the other datasets 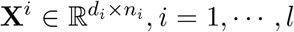 as the query datasets. We apply partial-GW to **X**^*i*^ and **Y** in the three steps above and obtain the probabilistic coupling matrices **T**^*i*^s of cells between **X**^*i*^s and **Y**, respectively, as the probabilistic cell-cell correspondence information.

We then align **X**^*i*^s and **Y** in a *d*_*e*_-dimensional common space, resulting in the new embeddings of 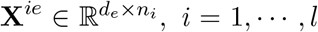, and 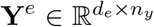. To preserve the local neighborhood relationship, we construct the graph Laplacian matrices 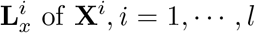, and **L**_*y*_ of **Y**, as other manifold learning algorithms have done [30–32]. Besides, we also introduce the rotation-invariant constraints and find the embeddings of cells by solving the optimization problem as

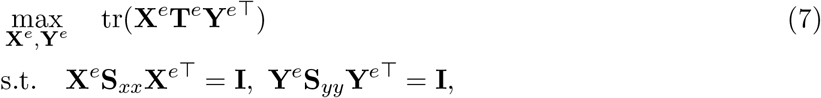

where 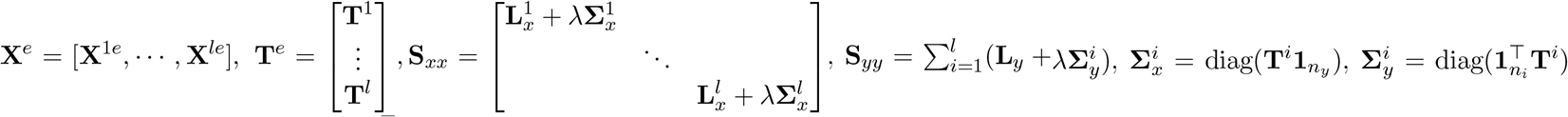 and tr(·) is the trace of matrix (see Supplementary Note 2 for details). We solve this optimization problem using the eigenvalue decomposition method as in [30, 33].

### Extension of partial-GW framework to incorporate prior information

Pamona can also incorporate existing prior information in the alignment, such as cell types or cell-cell correspondence, similar to the labeled graph matching problem [23]. Suppose each modality has its associated cell labels or correspondence, and the objective is to find an alignment that fits the prior information and data structure simultaneously. Let **M**_*ij*_ denote the disagreement between the *i*-th cell of dataset **X** and *j*-th cell of dataset **Y**. Then the alignment problem based on prior information can be formulated as

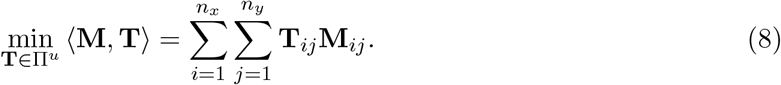

A natural way of unifying (2) and (8) to match both the data structure and the prior information is to minimize a convex combination [34]:

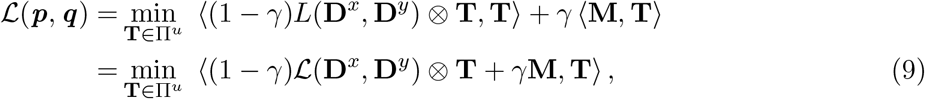

where *γ* is a non-negative regularization constant, and *γ* ∈ [0, 1], which represents a tradeoff between cost of individual matchings and faithfulness to the data structure.

However, being in different scales for different domain knowledge can affect integration performance considerably, e.g., a change in the scale of **M**. To overcome this problem, we introduce a more stable multiplication operator as

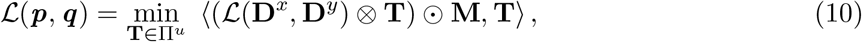

where ⊙ denotes Hadamard product which is the element-wise product taken on two matrices of the same dimensions. In implementation, we set **M**_*ij*_ = 1 − *γ* (0 < *γ* < 1) if cell *i* in dataset **X** and cell *j* in **Y** are in a correspondence relationship or the same cell type, and **M**_*ij*_ = 1 otherwise. A larger *γ* value gives more importance to the matching of prior information.

### Scree-Plot-Like estimation of the shared cell number *s*

The original partial-GW framework needs to specify the shared cell number *s*, i.e., the shared mass that needs to be transported. Nevertheless, in practice, *s* is generally unavailable. The user can choose *s* empirically. However, such an approach can either overestimate or underestimate *s*, leading to inaccurate alignment of the datasets.

The scree plot is used to determine the number of factors to retain in PCA [35]. In this study, we propose a SPL method that can accurately estimate shared cell number *ŝ* with errors < 10% of the true *s* in our tested data sets (Supplementary Fig. 14). The procedure of SPL is as follows.

Given a reference dataset **Y** with *n*_*y*_ cells and a query dataset **X** with *n*_*x*_ cells, we first give a rough guess of *s*, denoted as *s*^(*g*)^, which is smaller than *n*_*x*_, e.g., say in the range from 0.8 * *n*_*x*_ to 0.9 * *n*_*x*_. We apply Pamona with *s*^(*g*)^ and compute the corresponding cost matrix 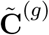 and the transport coupling matrix 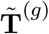. Then, we take out the last column of *T* ^(*g*)^ from 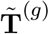, divide the interval between the minimum value and maximum value of *T* ^(*g*)^ into *d* equal spaced bins (small intervals), count the number of cells (the elements of *T* ^(*g*)^) falling into each bin, and plot the number of cells in each bin (Fig. 1b). Ideally, if query **X** has dataset-specific cells, they tend to be absorbed by the virtual point of **Y** with relatively high values in *T* ^(*g*)^. Thus, we can see a bump in the plot close to the side of maximum value of *T*^(*g*)^. We then detect the flat bin preceding the bump. To be specific, suppose each bin has *b*_*i*_ cells, where *i* ∈ 1, · · ·, *d*, and we want to find a bin with largest *i* satisfying the following

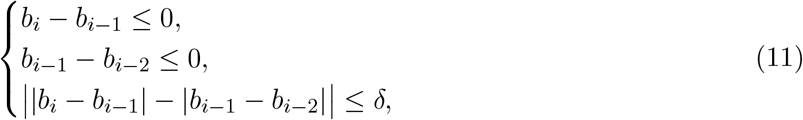

where |·| represent the absolute value, and *δ* = max{1, 0.005 × *n*_*x*_}. Once *i* is identified, we estimate *s* as 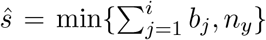. In this study, we applied SPL to estimate *ŝ* for all partial manifold alignment tasks.

### Method evaluations

We evaluate the single-cell multi-omics integration methods using three scores, (i) Label Transfer Accuracy, (ii) Alignment Score, and (iii) FOSCTTM, to measure the alignment accuracies. All three scores work on the basis of the common space (coordinate) of the integrated datasets.

Label Transfer Accuracy, which has been widely used in the transfer learning community and was adopted by UnionCom [9] and SCOT [10], is used to measure the ability to transfer labels of cells from one dataset to another in the common space. It is computed only using the shared cells and works when the cell label information (e.g., cell types, branches of cell trajectories) is available. We construct a *k*^*lta*^-nn classifier trained by using the reference dataset (e.g., Dataset **Y**) and compute the prediction accuracy of the shared cell labels on the query dataset (e.g., Dataset **X**) using the *k*^*lta*^-nn classifier. Label Transfer Accuracy is defined as the percentage of shared cells in the query dataset with correctly predicted labels, ranging from 0 to 100%. A higher Label Transfer Accuracy is indicative of better performance. In this study, we chose the parameter of the *k*^*lta*^-nn classifier as *k*^*lta*^ = max{10, 0.01 * *n*}, where *n* is the total number of cells in both query and reference datasets.

Alignment Score, which is the extension of the alignment score used in Seurat v2 [36], is applied to measure the ability to preserve both shared and dataset-specific structures in the common space. It works when the information of both shared and dataset-specific cells is available. First, for shared cells only, the procedure of Alignment Score is exactly the same as that in Seurat. Specifically, we randomly downsample the shared cells for larger datasets such that all datasets have the same number of shared cells as the dataset with the smallest shared cell number. Next we construct a *k*^*as*^ nearest-neighbor graph for all shared cells sampled based on their Euclidean distances in the common space. For each shared cell, we then compute how many of its *k*^*as*^ nearest-neighbors belong to the same dataset and compute the averaged value over all shared cells, denoted as 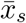. If the shared cells are well aligned, we would expect that the nearest neighbors of each shared cell to be uniformly shared across all datasets. For dataset-specific cells, we also construct a *k*^*as*^ nearest-neighbor graph of all shared and dataset-specific cells, based on their Euclidean distances in the common space, followed by computing how many of its *k*^*as*^ nearest-neighbors belong to the dataset-specific cells of the same dataset, and compute the averaged values over all dataset-specific cells, denoted as 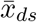. If the dataset-specific cells are well separated, we would expect that nearest neighbors of each dataset-specific cell to be all the dataset-specific cells in the same dataset. Suppose *l* +1 is the number of single-cell multi-omics datasets. Then the Alignment Score is defined as:

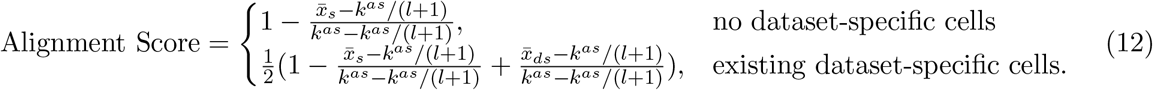

A higher Alignment Score is indicative of better performance. In this study, we chose the parameter of *k*^*as*^ = max{10, 0.01 * *n*}, where *n* is the total number of the shared cells.

The FOSCTTM, or fraction of samples closer than the true match, score was introduced by MMD-MA [7] to measure the preservation of cell-cell relationships in the common space when the information of cell-cell correspondence across datasets is available. We compute the Euclidean distances between a fixed sample point from one dataset and all the data points in the other dataset. Next, we compute the fraction of those distances that are closer to the sample point than the distance to the true matched point (correspondence cell). The FOSCTTM score is defined as the averaged value of the fractions over all the samples. For perfect alignment, all samples would be closest to their true match, yielding a FOSCTTM score of zero. Therefore, a lower FOSCTTM score is indicative of better performance.

### Data and data preprocessing

Our Simulation 1 data set was first simulated by SCOT [10]. It embedded a bifurcated tree into two datasets: dataset **X** has 300 cells and 1000 features; dataset **Y** has 300 cells and 2000 features, respectively.

Our Simulation 2 data set was first simulated by UnionCom [9]. It embedded a bifurcated tree into dataset **X**, which has 200 cells and 1000 features, and embedded a trifurcated tree into dataset **Y**, which has 250 cells and 500 features. Dataset **Y** has a dataset-specific branch.

The sc-GEM data set of samples from human cells undergoing reprogramming to iPSCs was generated by [17]. We utilized the preprocessed sc-GEM data obtained from MATCHER [6] and the cell type annotation obtained from Cheow et al. [17]. It contains a DNA methylation dataset with 177 cells and 27 features and a gene expression dataset with 177 cells and 34 features (genes), respectively.

The scNMT-seq data set of mouse gastrulation samples collected at 4 time stages was generated by [18]. It contains 3 datasets of RNA, DNA methylation and chromatin accessibility. We followed the data analysis pipeline of scNMT-seq [18] and chose samples annotated as “E4.5 Epiblast”, “E5.5 Epiblast”, “E6.5 Epiblast”, “E6.5 Primitive Streak”, “E6.5 Mesoderm”, “E7.5 Epiblast”, “E7.5 Primitive Streak”, “E7.5 Ectoderm”, “E7.5 Endoderm”, and “E7.5 Mesoderm” for the integration, resulting in 2147 cells with 3473 genes in RNA, 647 cells with 10000 features in chromatin accessibility, and 725 cells with 5000 features in DNA methylation, respectively. We further filtered samples of the RNA dataset and selected out cells from the first batch of scRNA-seq of mouse embryos E4.5 to E7.5 in Gene Expression Omnibus accession GSE121708, resulting in 597 cells retained in the RNA dataset. Finally, we performed PCA dimensionality reduction for each of the three modalities and retained the top 30 components. The preprocessed scNMT-seq data set contains a gene expression dataset with 597 cells and 30 features, a chromatin accessibility dataset with 647 cells and 30 features, and a DNA methylation dataset with 725 cells and 30 features, respectively.

The SNARE-seq data set of samples, derived from a mixture of BJ, H1, K562, and GM12878 cell lines, was generated by [19] and utilized by SCOT [10]. We followed the data preprocessing procedure of SCOT as follows: We performed dimensionality reduction of the scATAC-seq dataset using cisTopic [37] and applied PCA for dimensionality reduction of the scRNA-seq dataset. The preprocessed SNARE-seq data set contains an scRNA-seq dataset with 1047 cells and 10 features and an scATAC-seq dataset with 1047 cells and 19 features, respectively.

The PBMC data set of samples of human peripheral blood mononuclear cells was generated by 10X Genomics. The preprocessed PBMC data set, which was obtained from MAESTRO [20], contains an scATAC-seq dataset with 1919 cells and 50 features and an scRNA-seq dataset with 1985 cells and 50 features, respectively. We followed the cell type annotation functions “ATACAnnotationCelltype.R” and “RNAAnnotationCelltype.R” provided by MAESTRO to annotate scATAC-seq and scRNA-seq data, respectively, resulting in cell types “CD8Tex”, “Monocytes”, “NaiveCD4Tcells” and “RestNK” for both scATAC-seq and scRNA-seq datasets, “Tfh” and “MemoryBcells” for the scATAC-seq dataset only, and “CD8Tcells”, “RestMemCD4Tcells”, “RestpDCs” and “NaiveBcells” for the scRNA-seq dataset only. We normalized the features across samples within each of the datasets before alignment using z-score method.

### Computational complexity and time

Memory complexity of Pamona is *O*(*n*^2^), where *n* is the number of samples. Time complexity of Pamona is *O*(*kn*^3^) with *k* iterations, which is similar to GW-based methods (e.g. SCOT in Supplementary Fig. 9d). Our experimental environment includes Intel Core CPU i9-9980XE 3.0GHz, 128GB DDR4 memory and NVIDIA GPU TITAN RTX. For data sets of Simulation 1, Simulation 2, sc-GEM and scNMT-seq, running time of Pamona is less than 20 seconds. For PBMC data set, it is around 2 minutes, and for SNARE-seq data set, it is around 7 minutes.

## Data availability

The simulated data is available at https://noble.gs.washington.edu/proj/mmd-ma.

The sc-GEM data is available at https://github.com/jw156605/MATCHER/tree/master/pymatcher/data. The scNMT-seq data is available at https://www.ncbi.nlm.nih.gov/geo/query/acc.cgi?acc=GSE121708 and ftp://ftp.ebi.ac.uk/pub/databases/scnmt_gastrulation/scnmt_gastrulation.tar.gz.

The SNARE-seq data is available at http://www.ncbi.nlm.nih.gov/geo/query/acc.cgi?acc=GSE126074 (preprocessed data available at https://github.com/rsinghlab/SCOT/tree/master/data).

The PBMC data is available at https://github.com/liulab-dfci/MAESTRO/tree/master/data.

## Code availability

Pamona software is available at https://github.com/caokai1073/Pamona. The implementation of the partial GW framework is based on the POT: Python Optimal Transport toolbox [38].

## Acknowledgements

This work was supported by the National Key R&D Program of China under Grant 2019YFA0709501, NSFC grants (Nos. 61733018, 12071466), LSC of CAS.

