## Supplementary Methods and Figures for "Manifold alignment for heterogeneous single-cell multi-omics data integration using Pamona"

#### Supplementary Note 1: Pamona algorithm

**Algorithm 1** Pamona algorithm

Input: Query datasets $\mathbf{X}^i,\ i = 1, \cdots, l$, reference dataset $\mathbf{Y}$ and shared cell number $s$;

Initialize: $\tilde{\boldsymbol{p}}, \tilde{\boldsymbol{q}}, \epsilon, \mathbf{T}^i, i = 1 \cdots, l$;
**for** $i = 0$ to $l$ **do**
  $\mathbf{X} = \mathbf{X}^i$, $\mathbf{T}^{(0)} = \mathbf{T}^i$;
  Construct geodesic distance matrices $\mathbf{D}^x$ and $\mathbf{D}^y$;
  $k = 0$, err $= \epsilon + 1$;
  **while** $k \leq K$ and err $> \epsilon$ **do**
    Compute the cost matrix $\mathbf{C}^{(k)}$ as Eq. (3);
    If disagreement matrix $\mathbf{M}$ of dataset $\mathbf{X}$ and $\mathbf{Y}$ is available, $\mathbf{C}^{(k)} \leftarrow \mathbf{C}^{(k)} \odot \mathbf{M}$;
    Construct the augmented cost matrix $\tilde{\mathbf{C}}^{(k)}$ as Eq. (4);
    Compute $\tilde{\mathbf{T}}^{(k+1)}$ in Eq. (5) by Sinkhorn iterations [3,6];
    Remove the last column and row of $\tilde{\mathbf{T}}^{(k+1)}$ to obtain $\mathbf{T}^{(k+1)}$;
    err $= \|\mathbf{T}^{(k+1)} - \mathbf{T}^{(k)}\|_F$;
    $k = k + 1$;
  $\mathbf{T}^i = \mathbf{T}^{(k)}$;
Construct graph Laplacian matrices $\mathbf{L}_x^i,\ i = 1, \cdots, l$, and $\mathbf{L}_y$;
Compute $\mathbf{\Sigma}_x^i = \mathrm{diag}(\mathbf{T}^i \mathbf{1}_{n_y}),\ \mathbf{\Sigma}_y^i = \mathrm{diag}(\mathbf{1}_{n_x}^\top \mathbf{T}^i),\ i = 1, \cdots, l,\ \mathbf{S}_{xx},\ \mathbf{S}_{yy}$ and $\mathbf{T}^e$ in Eq. (7);
$\mathbf{H} = \mathbf{S}_{xx}^{-1/2} \mathbf{T}^e \mathbf{S}_{yy}^{-1/2}$;
Eigenvalue decomposition $\mathbf{H} = \mathbf{U}\mathbf{\Sigma}\mathbf{V}^\top$;
$\mathbf{X}^e = \mathbf{S}_{xx}^{-1/2}\mathbf{U}$, $\mathbf{Y}^e = \mathbf{S}_{yy}^{-1/2}\mathbf{V}$;
**return** $\mathbf{T}^e, \mathbf{X}^e, \mathbf{Y}^e$

#### Supplementary Note 2: Detailed information of step 4 of Pamona

For the case that we have $l + 1$ ($l \geq 1$) multiple datasets, we fix a dataset $\mathbf{Y} \in \mathbb{R}^{d_y \times n_y}$ as the reference dataset, and the other datasets $\mathbf{X}^i \in \mathbb{R}^{d_i \times n_i}, i = 1, \cdots, l$ as query datasets. We apply partial-GW to $\mathbf{X}^i$ and $\mathbf{Y}$, and obtain the probabilistic coupling matrices $\mathbf{T}^i$s of cells between $\mathbf{X}^i$s and $\mathbf{Y}$ as the correspondence information of cells. To preserve the local neighborhood relationship, we construct the graph Laplacian matrices $\mathbf{L}_x^i, i = 1, \cdots, l$ of dataset $\mathbf{X}^i$, and $\mathbf{L}_y$ of dataset $\mathbf{Y}$, as other manifold learning algorithms have done [1,2,7]. Therefore, we define the geometry preserving term as

$$E_g = \sum_{i=1}^{l} (\mathrm{tr}(\mathbf{X}^{ie} \mathbf{L}_x^i \mathbf{X}^{ie\top}) + \mathrm{tr}(\mathbf{Y}^e \mathbf{L}_y \mathbf{Y}^{e\top})),$$

where $\mathbf{X}^{ie} \in \mathbb{R}^{d_e \times n_i}$ and $\mathbf{Y}^e \in \mathbb{R}^{d_e \times n_y}$ are new embeddings of $\mathbf{X}^i$ and $\mathbf{Y}$ in a $d_e$-dimensional space, and $\mathrm{tr}(\cdot)$ is the trace of matrix.

To partial align all datasets in a common embedded space, we utilize the coupling matrices

$\mathbf{T}^i, i = 1, \cdots, l$, and formulate the feature alignment term as:

$$E_a = \sum_{i=1}^{l}\sum_{j=1}^{n_i}\sum_{k=1}^{n_y} \|\boldsymbol{x}_j^{ie} - \boldsymbol{y}_k^e\|^2 \mathbf{T}_{jk}^i = \sum_{i=1}^{l} \mathrm{tr}(\mathbf{X}^{ie}\mathbf{\Sigma}_x^i \mathbf{X}^{ie\top} + \mathbf{Y}^e \mathbf{\Sigma}_y^i \mathbf{Y}^{e\top} - 2\mathbf{X}^{ie}\mathbf{T}^i\mathbf{Y}^{e\top}),$$

where $\boldsymbol{x}_i^{ie}(\boldsymbol{y}_k^e)$ represents the $j(k)$-th column of matrix $\mathbf{X}^{ie}(\mathbf{Y}^e)$, and $\mathbf{\Sigma}_x^i = \mathrm{diag}(\mathbf{T}^i\mathbf{1}_{n_y})$, $\mathbf{\Sigma}_y^i = \mathrm{diag}(\mathbf{1}_{n_x}^\top \mathbf{T}^i)$. Consequently, we formulate the problem as the following optimization problem:

$$\begin{aligned}\min_{\substack{\mathbf{Y}^e, \mathbf{X}^{ie}, \\ i=1\cdots,l}} \quad & \mathcal{J}(\mathbf{X}^e, \mathbf{Y}^e) = E_g + \lambda E_a \\ & = \sum_{i=1}^{l} \mathrm{tr}(\mathbf{X}^{ie}(\mathbf{L}_x^i + \lambda\mathbf{\Sigma}_x^i)\mathbf{X}^{ie\top} + \mathbf{Y}^e(\mathbf{L}_y + \lambda\mathbf{\Sigma}_y^i)\mathbf{Y}^{e\top} - 2\lambda\mathbf{X}^{ie}\mathbf{T}^i\mathbf{Y}^{e\top}),\end{aligned}$$

where $\lambda$ is a balance parameter. We introduce the rotation-invariant constraints, and reformulate the optimization problem as:

$$\begin{aligned}\max_{\mathbf{X}^e, \mathbf{Y}^e} \quad & \mathrm{tr}(\mathbf{X}^e\mathbf{T}^e\mathbf{Y}^{e\top}) \\ \text{s.t.} \quad & \mathbf{X}^e\mathbf{S}_{xx}\mathbf{X}^{e\top} = \mathbf{I}, \ \mathbf{Y}^e\mathbf{S}_{yy}\mathbf{Y}^{e\top} = \mathbf{I},\end{aligned}$$

where $\mathbf{X}^e = [\mathbf{X}^{1e}, \cdots, \mathbf{X}^{le}]$, $\mathbf{T}^e = \begin{bmatrix}\mathbf{T}^1 \\ \vdots \\ \mathbf{T}^l\end{bmatrix}$, $\mathbf{S}_{xx} = \begin{bmatrix}\mathbf{L}_x^1 + \lambda\mathbf{\Sigma}_x^1 & & \\ & \ddots & \\ & & \mathbf{L}_x^l + \lambda\mathbf{\Sigma}_x^l\end{bmatrix}$ and $\mathbf{S}_{yy} = \sum_{i=1}^{l}(\mathbf{L}_y + \lambda\mathbf{\Sigma}_y^i)$. We solve this optimization problem using the eigenvalue decomposition method as in [2,4].

#### Supplementary Note 3: Hyperparameter tuning

Pamona has the two same parameters as SCOT, and has two unique parameters in the manifold alignment: the number of neighborhoods $k$ in the $k$-nn graph is typically set to $k = \{10, 30, 50\}$; the regularization parameter $\epsilon$ is typically set to $\epsilon = \{0.0005, 0.001\}$; the trade-off parameter $\lambda$ is typically set to $\lambda = \{1, 10\}$; the dimensionality $d_e$ of common space is typically set to $d_e = \{5, 30\}$.

For UnionCom, we searched the trade-off parameter $\beta \in \{0.1, 1, 10\}$, $epoch_{\mathrm{DNN}} \in \{200, 500\}$, $\epsilon \in \{0.0001, 0.001\}$ for best performance. Other parameters of UnionCom were set to default values. For SCOT, because it shares the same two parameters $\epsilon$ and $k$ with Pamona, we set the same values as Pamona. For MMD-MA, we searched trade-off parameters $\lambda_1 \in \{1e-03, 1e-04, 1e-05, 1e-06, 1e-07\}$ and $\lambda_2 \in \{1e-03, 1e-04, 1e-05, 1e-06, 1e-07\}$ for best performance, and set the dimensionality of the common low-dimensional space $p = 5$ as default. Four Seurat v3, we set parameters to default values for all experiments.

### Supplementary Figures

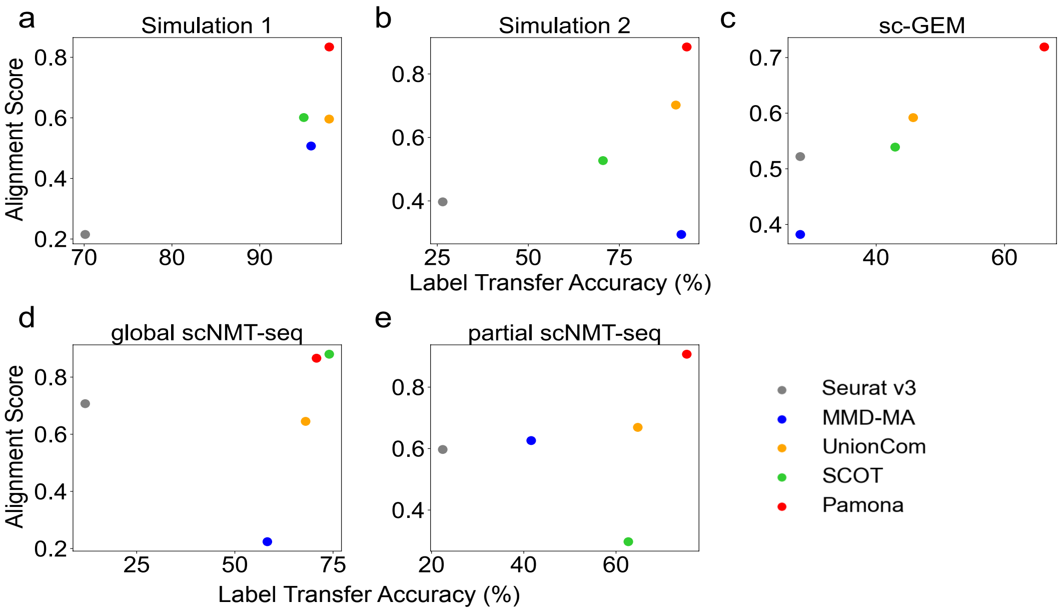


**Supplementary Figure 1.** Comparison of performance of Seurat v3, MMD-MA, UnionCom, SCOT and Pamona in Label Transfer Accuracy and Alignment score for partial manifold alignment tasks on Simulation 1, Simulation 2, sc-GEM, and scNMT-seq data sets, and a global manifold alignment task on scNMT-seq data set.

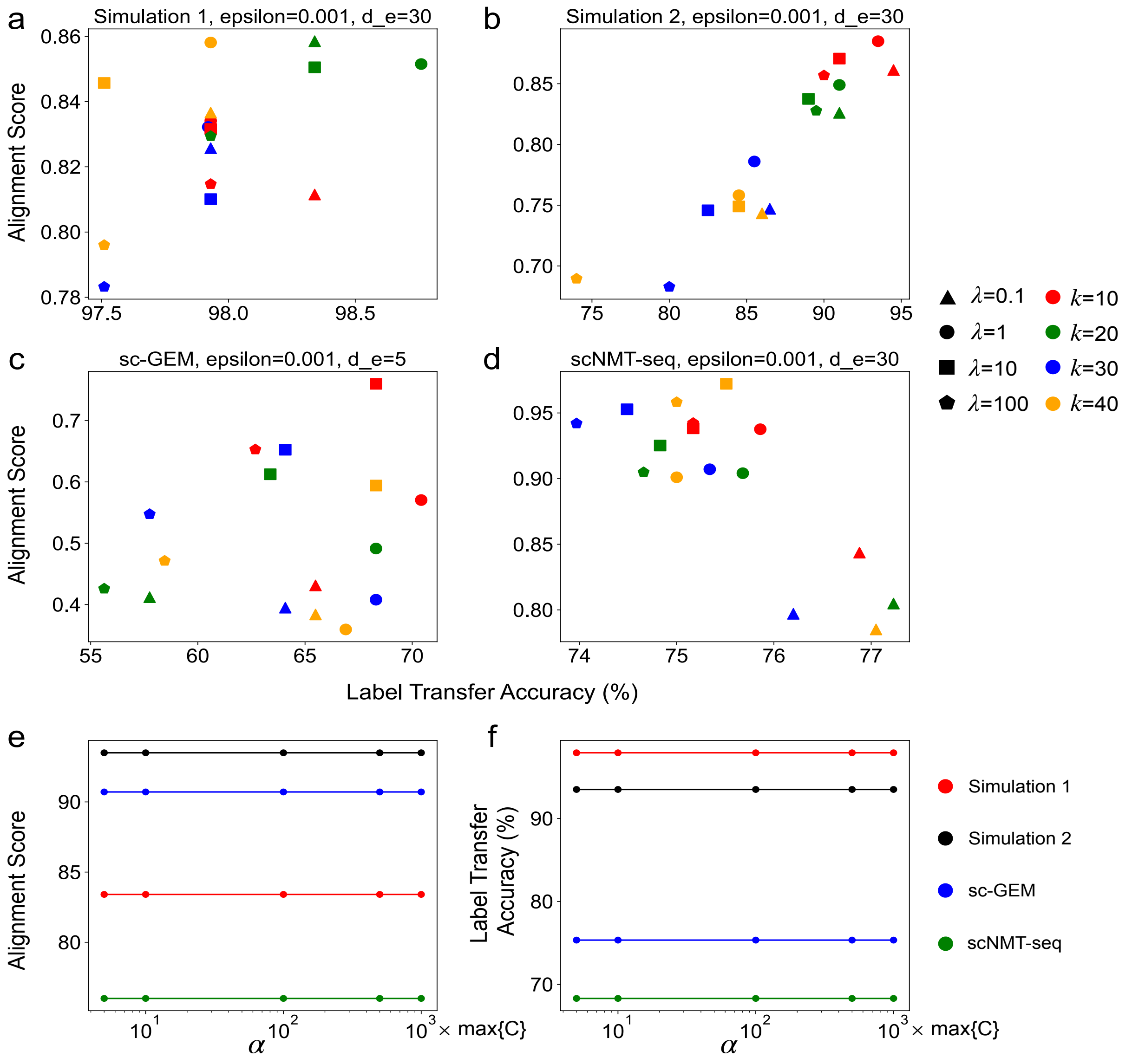


**Supplementary Figure 2.** Robustness of Pamona to parameter choices. (a-d) The trade-off parameter $\lambda$, the number of neighborhoods $k$. (e-f) The alignment cost of $\alpha$ within virtual cells.

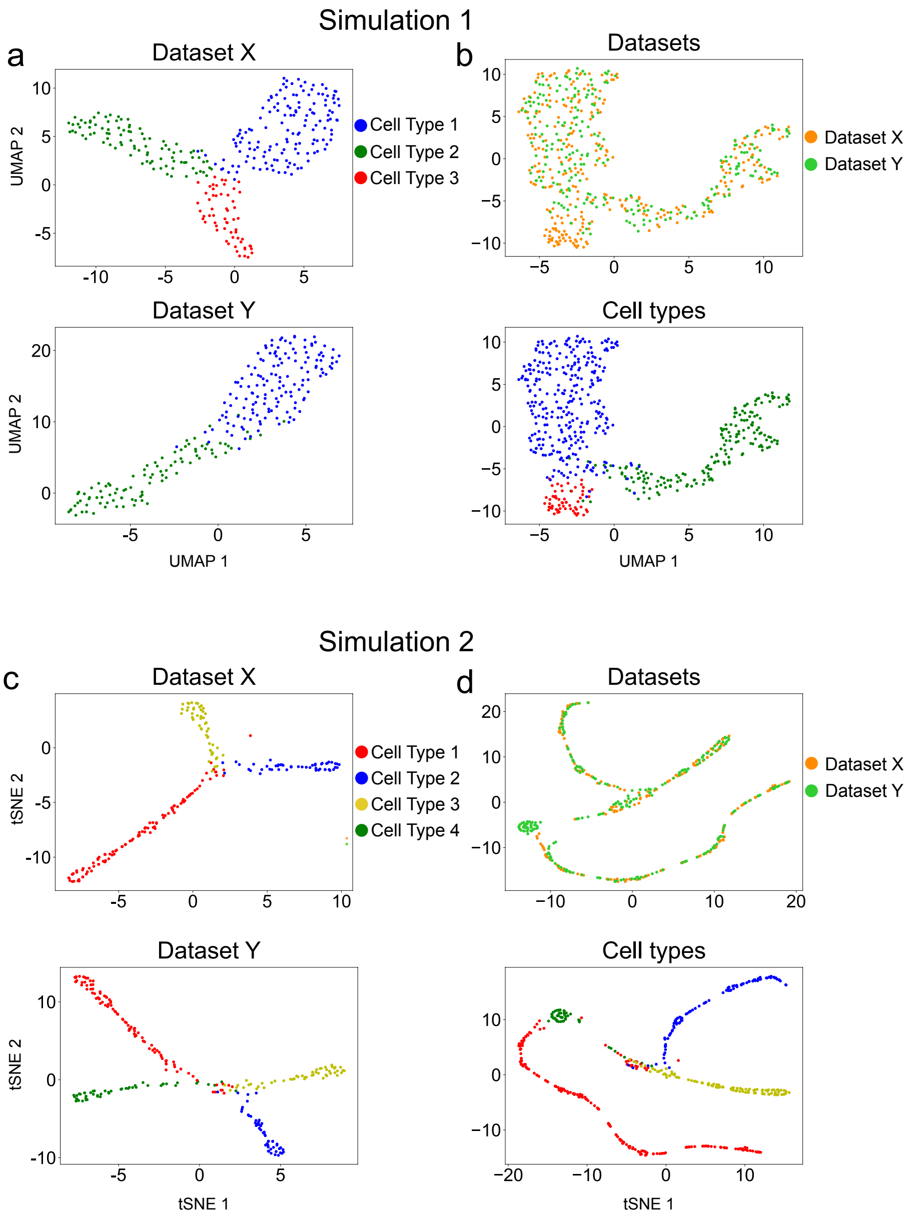


**Supplementary Figure 3.** Pamona integrated two simulated data sets in partial manifold alignment tasks. (a) Visualizations of the datasets **X** (upper panel) and **Y** (lower panel) in Simulation 1 separately using UMAP before alignment. (b) Visualizations of the common space of the partial alignment of the two aligned datasets in Simulation 1 by Pamona using UMAP: upper panel: cells are colored according to their corresponding datasets; lower panel: cells are colored according to their corresponding types. (c) Visualizations of the datasets **X** (upper panel) and **Y** (lower panel) in Simulation 2 separately using tSNE [5] before alignment. (d) Visualizations of the common space of the partial alignment of the two aligned datasets in Simulation 2 by Pamona using tSNE.

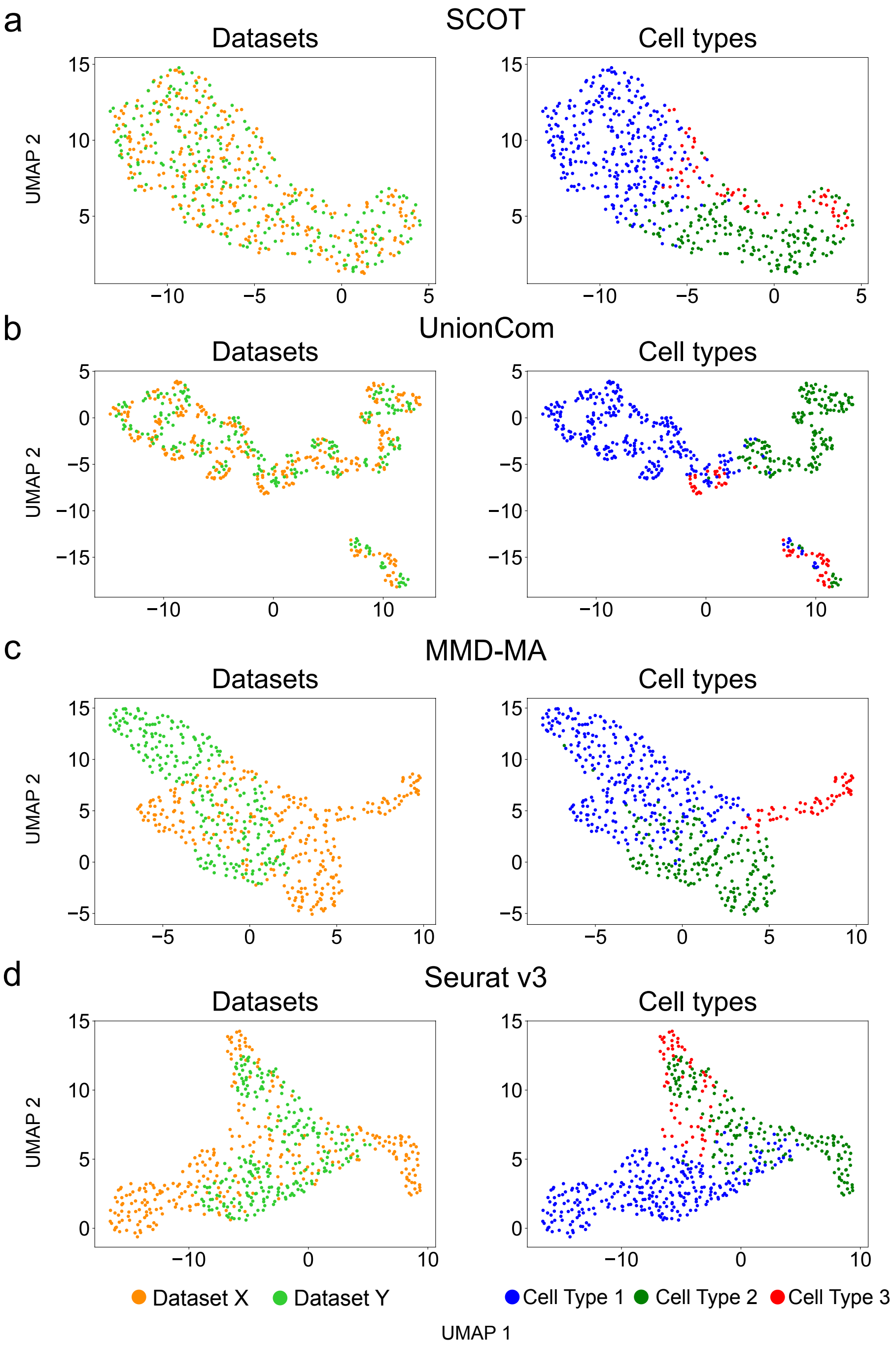


**Supplementary Figure 4.** Partial alignment results by SCOT (a), UnionCom (b), MMD-MA (c) and Seurat v3 (d) on Simulation 1.

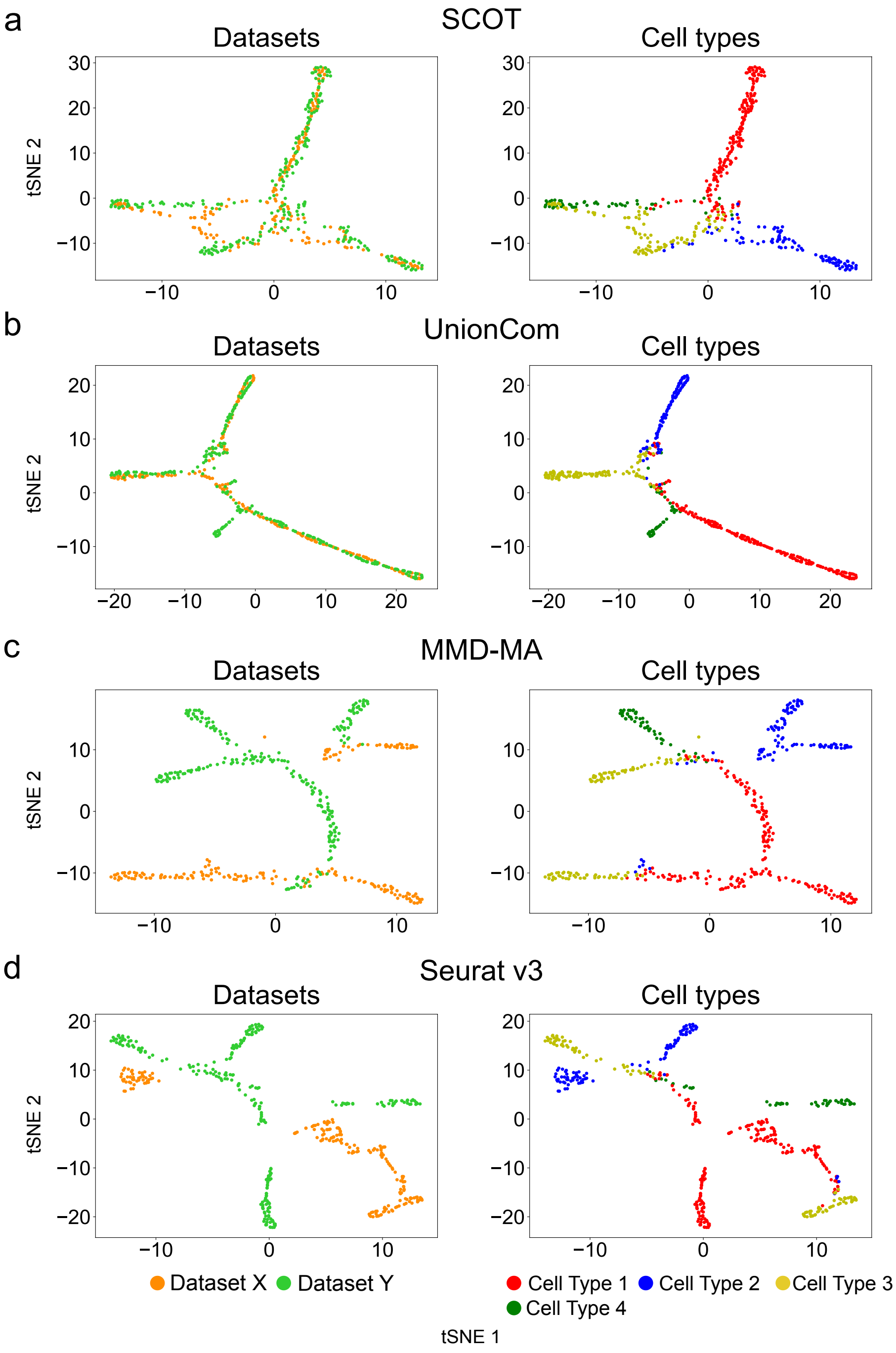


**Supplementary Figure 5.** Partial alignment results by SCOT (a), UnionCom (b), MMD-MA (c) and Seurat v3 (d) on Simulation 2.

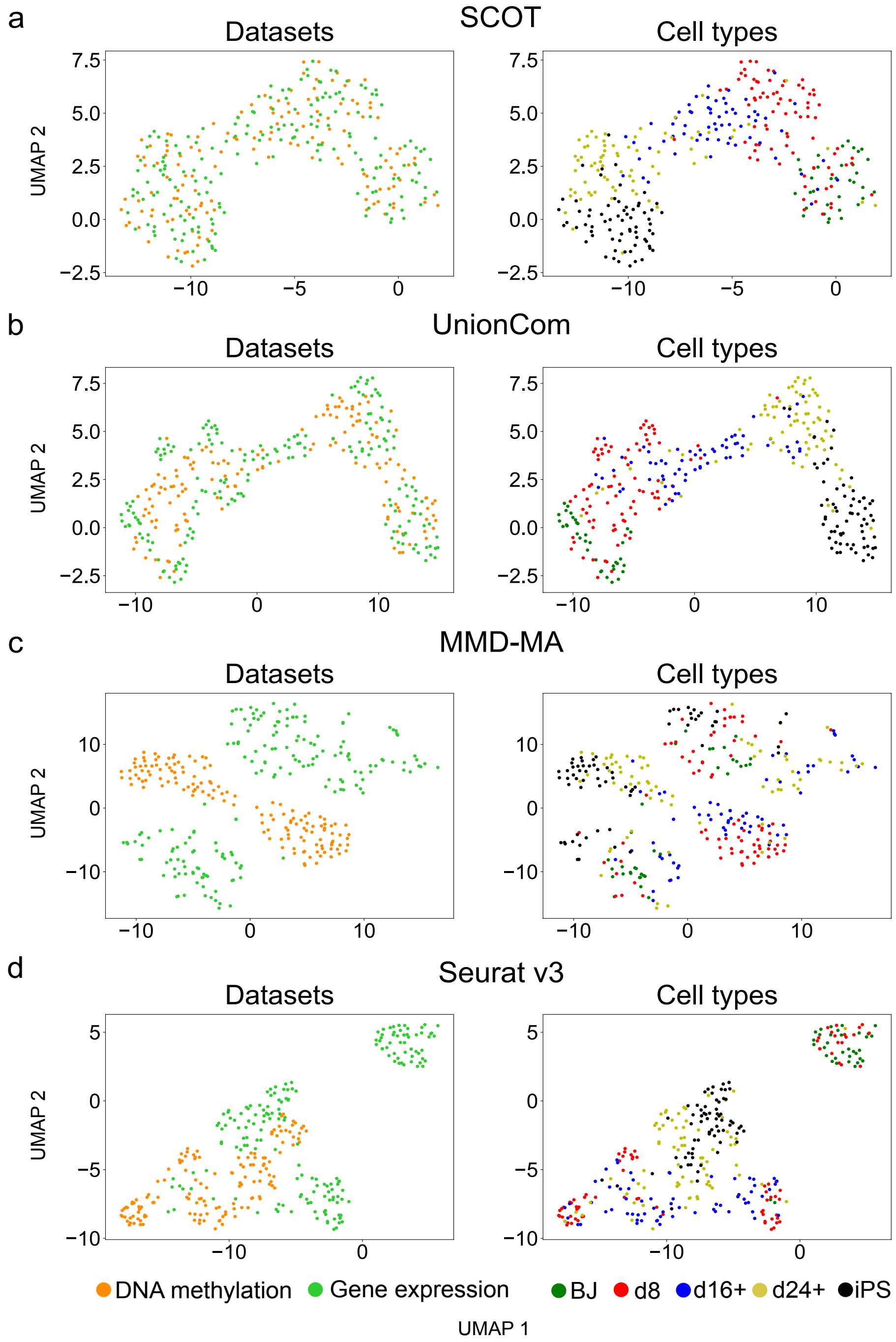


**Supplementary Figure 6.** Partial alignment results by SCOT (a), UnionCom (b), MMD-MA (c) and Seurat v3 (d) on sc-GEM data set.

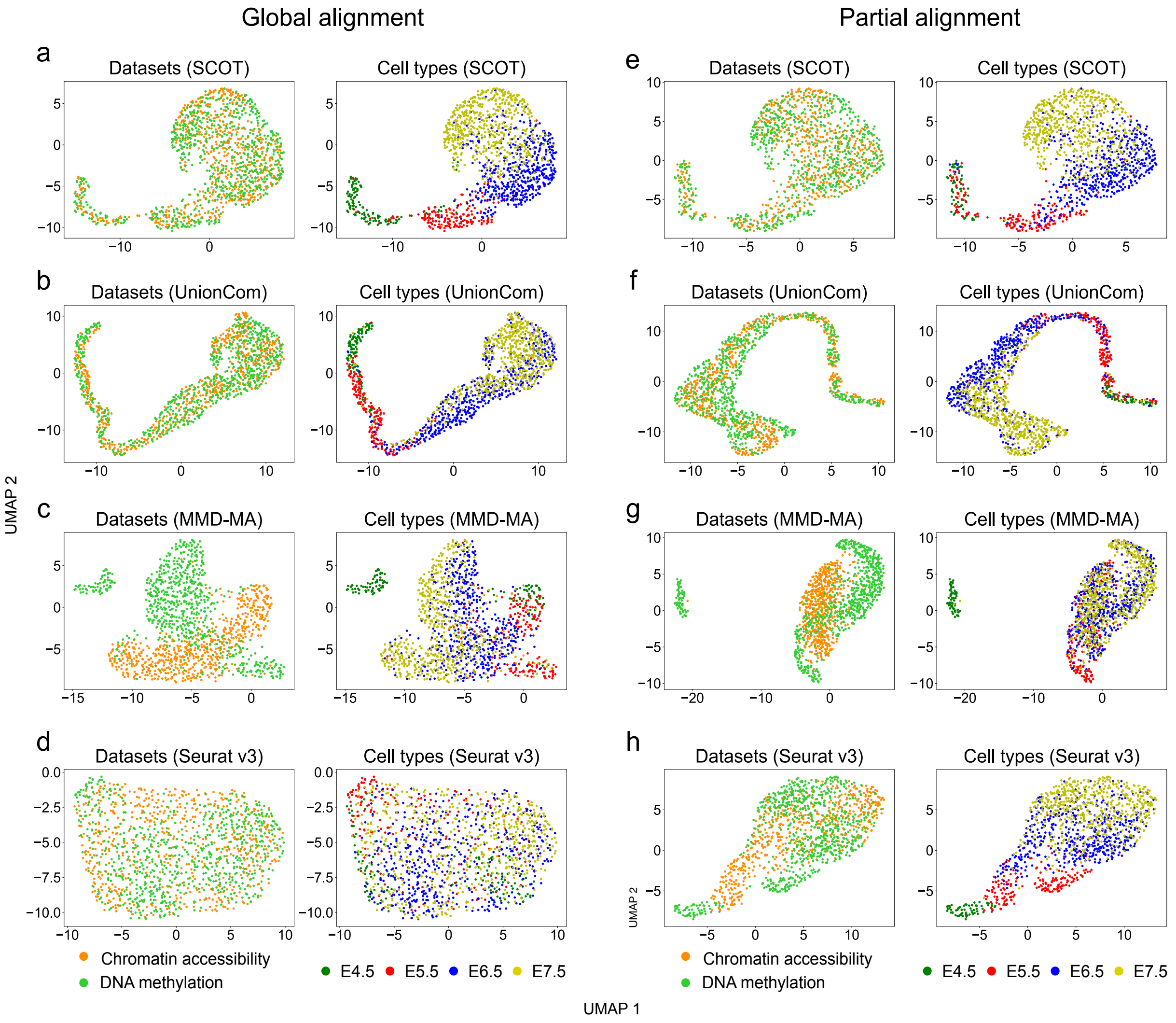


**Supplementary Figure 7.** Global and partial alignment results by SCOT (a - global, e - partial), UnionCom (b - global, f - partial), MMD-MA (c - global, g - partial) and Seurat v3 (d - global, h - partial) on scNMT-seq data set.

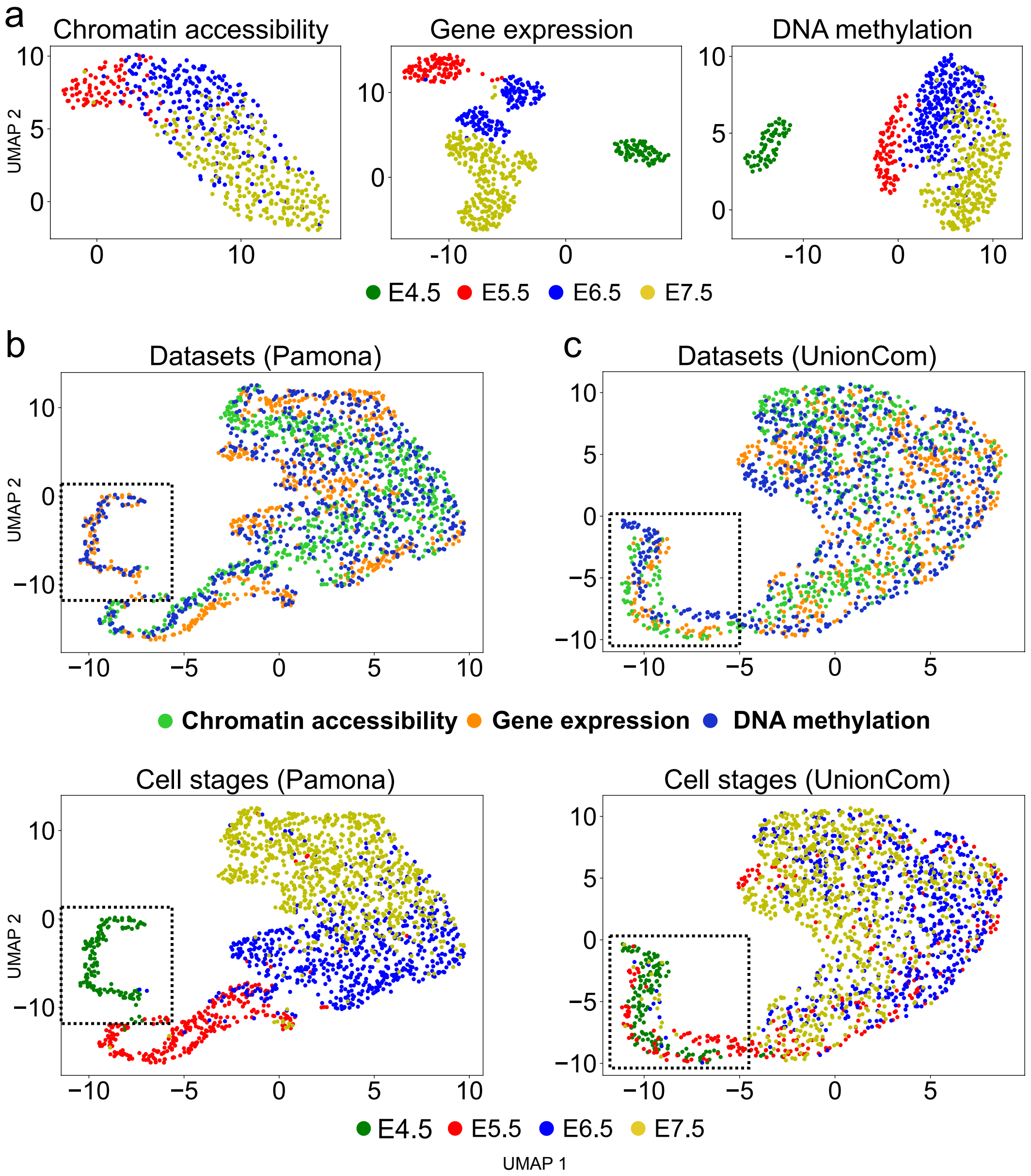


**Supplementary Figure 8.** Pamona accurately integrated partial scNMT-seq datasets of chromatin accessibility, gene expression and DNA methylation. (a) Visualizations of the chromatin accessibility, gene expression and DNA methylation datasets separately using UMAP before alignment. (b) Visualization of the common space of the partial alignment of the three aligned datsets by Pamona using UMAP (upper panel: points are colored according to their corresponding datasets; lower panel: points are colored according to their corresponding types). (c) Visualization of the common space of the partial alignment of the three aligned datsets by UnionCom using UMAP.

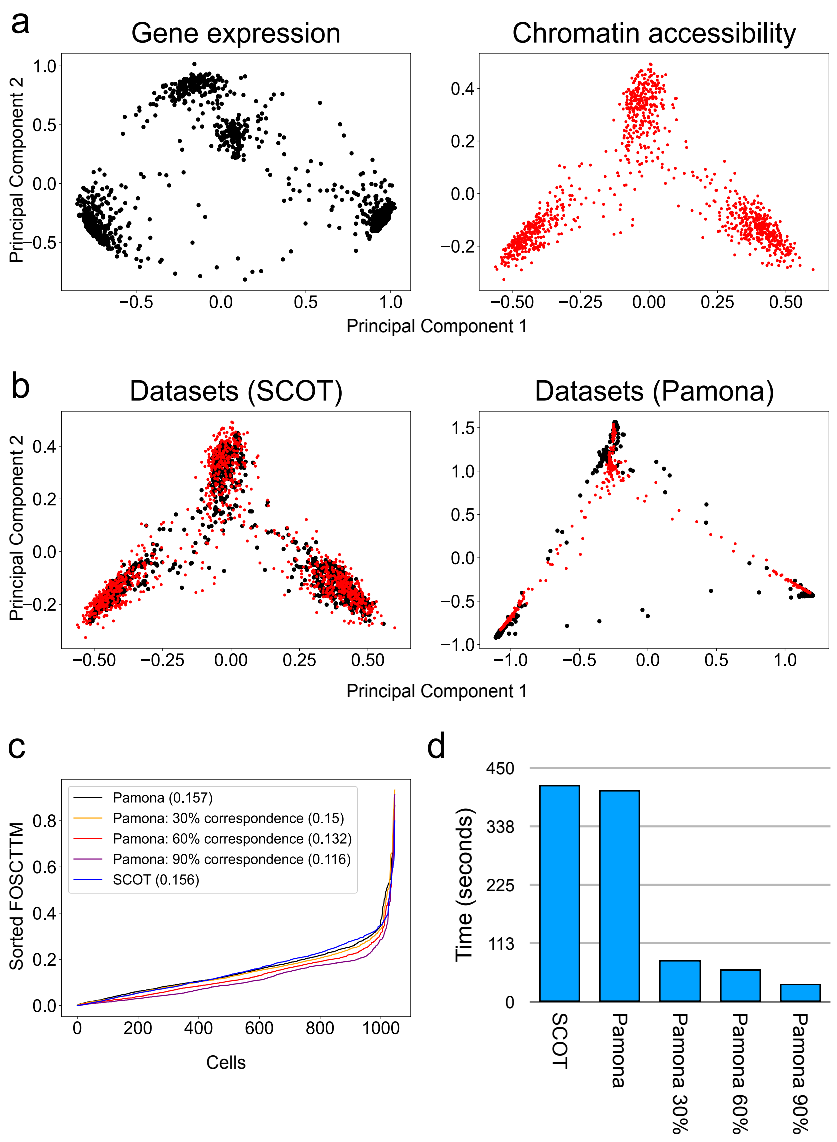


**Supplementary Figure 9.** Pamona accurately integrated SNARE-seq datasets of gene expression and chromatin accessibility by incorporating partial cell-cell correspondence information. (a) Visualizations of the gene expression and chromatin accessibility datasets separately using PCA before alignment. (b) Visualizations of the common space of the global alignment of the two datasets by Pamona and SCOT without cell-cell correspondence using PCA (points are colored according to their corresponding datasets). By incorporating prior information, Pamona improved the performance of cell-cell correspondence perservation measured by the FOSCTTM score (c) , and showed up to 10-fold acceleration of computational speed (d).

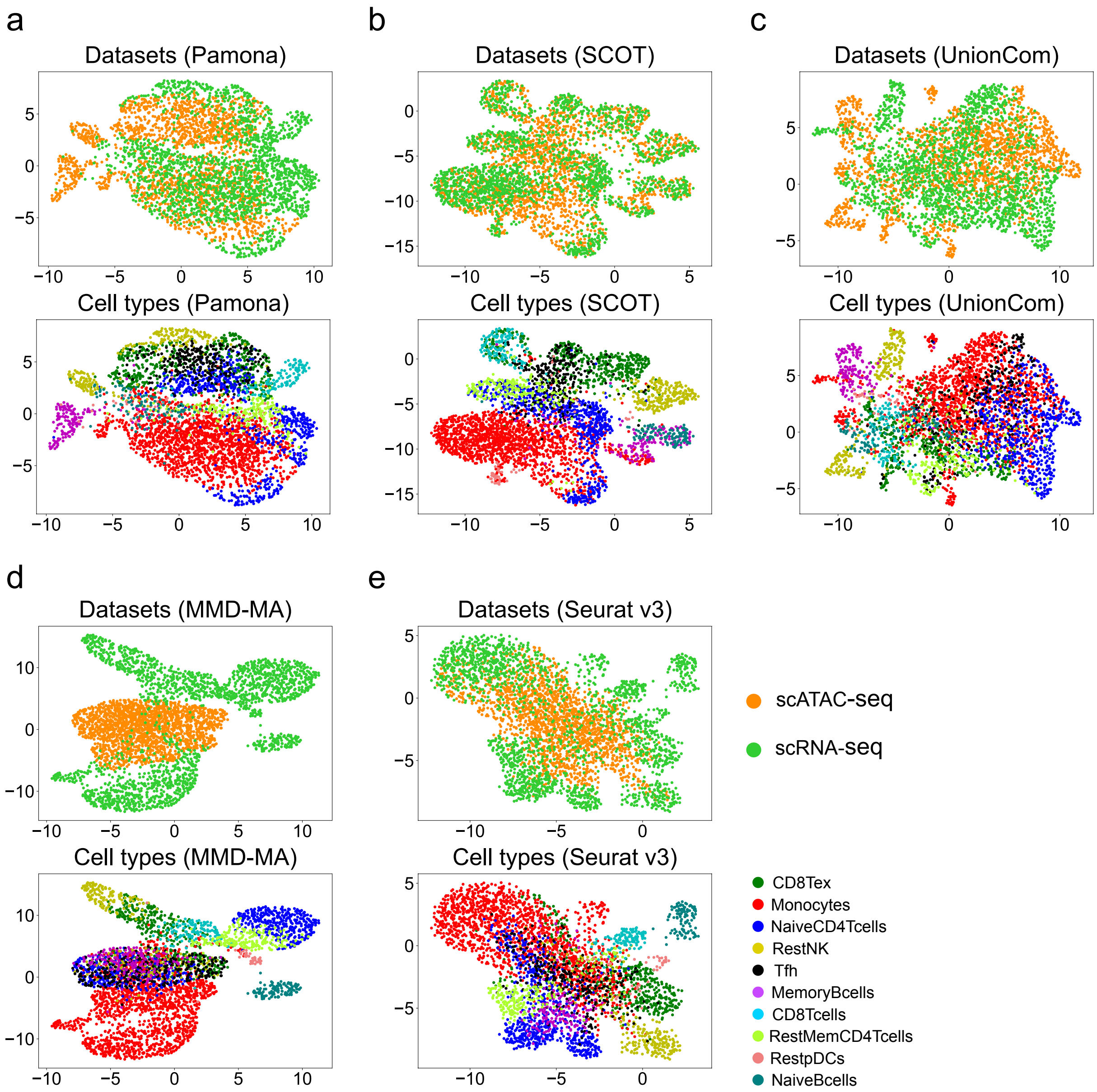


**Supplementary Figure 10.** Partial alignment results by Pamona (a), SCOT (b), UnionCom (c), MMD-MA(d) and Seurat v3 (e) on PBMC scATAC-seq and scRNA-seq data when no cell type information is incorporated.

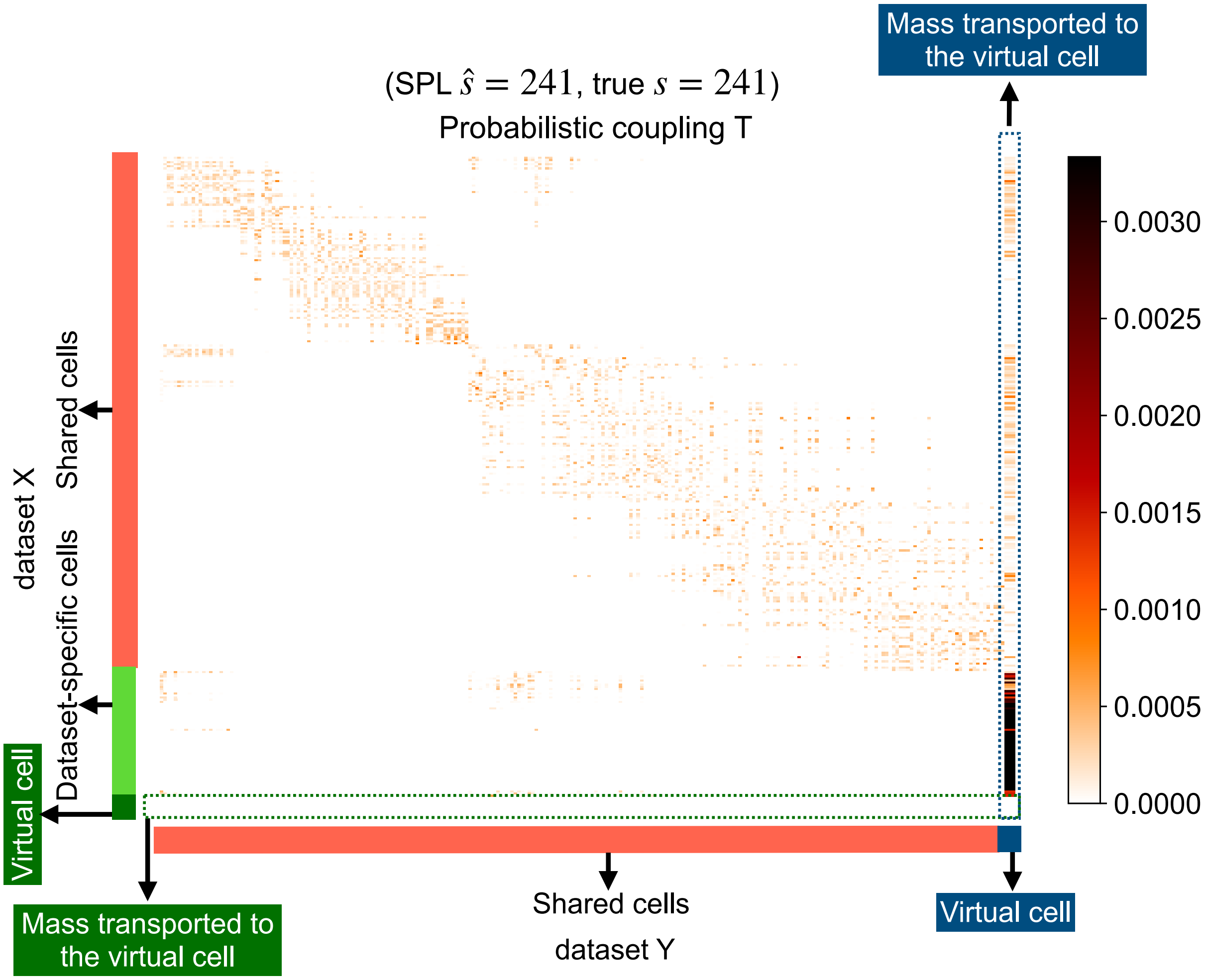


**Supplementary Figure 11.** Probabilistic coupling **T** of Pamona on Simualtion 1.

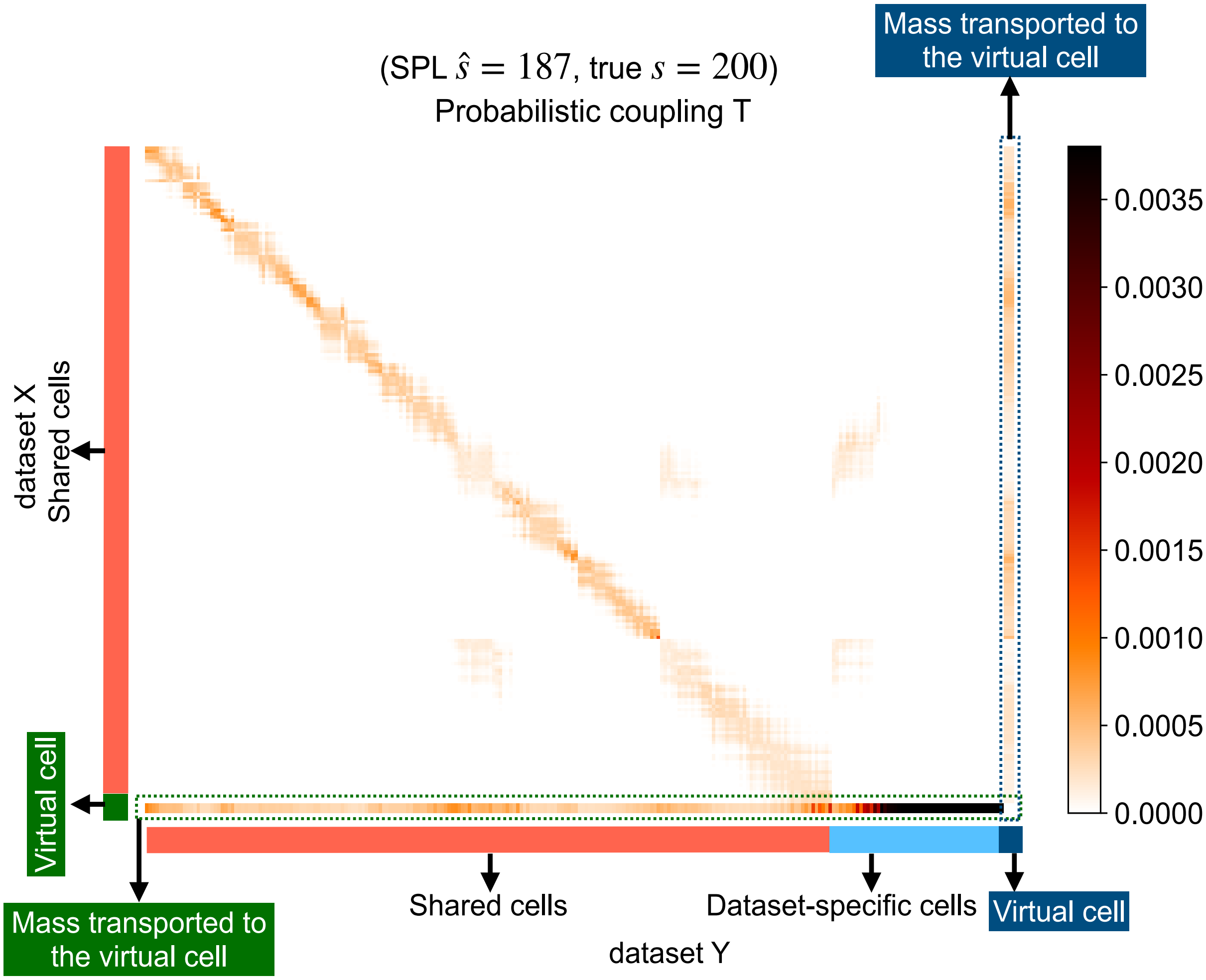


**Supplementary Figure 12.** Probabilistic coupling **T** of Pamona on Simualtion 2.

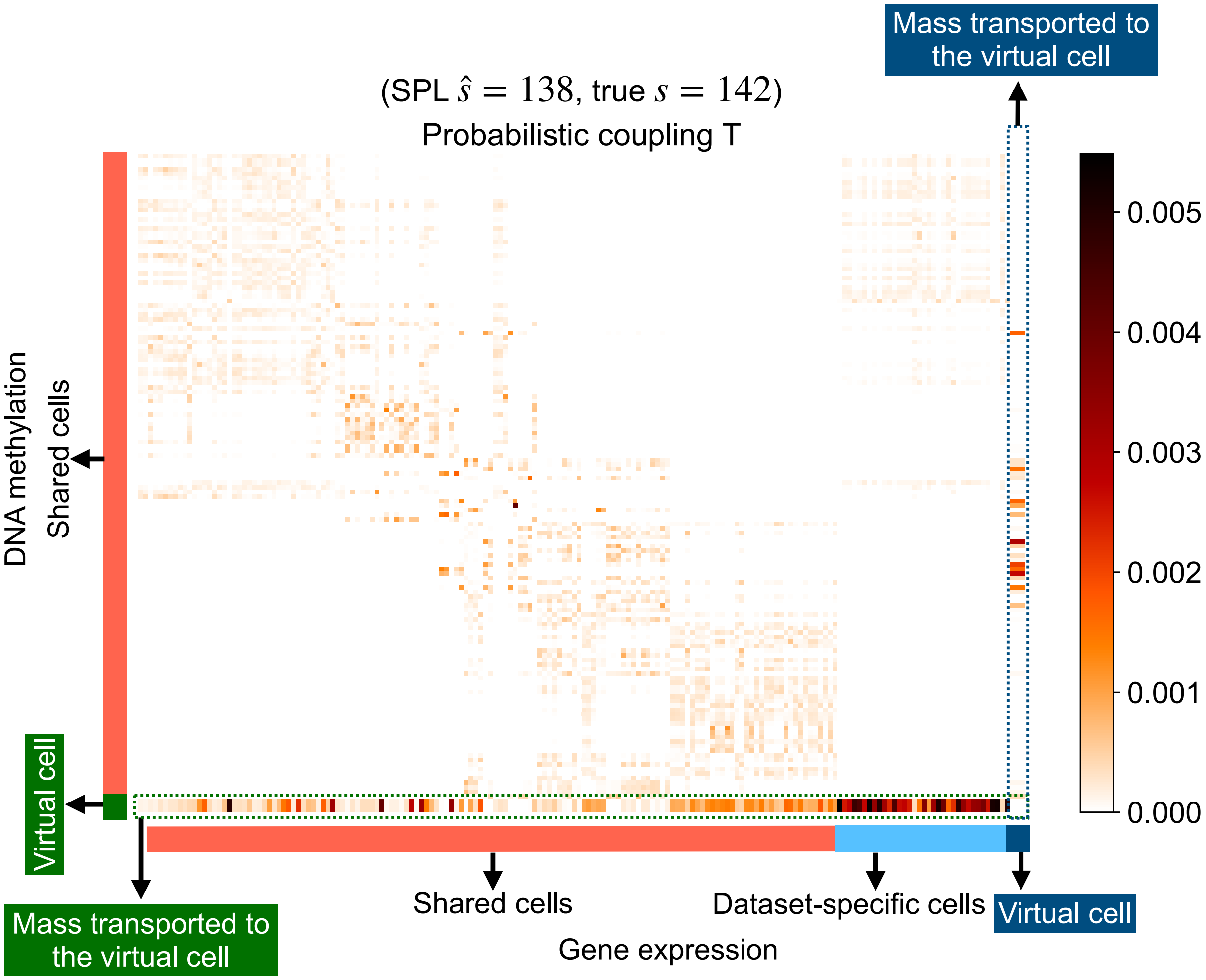


**Supplementary Figure 13.** Probabilistic coupling **T** of Pamona on scGEM data set.

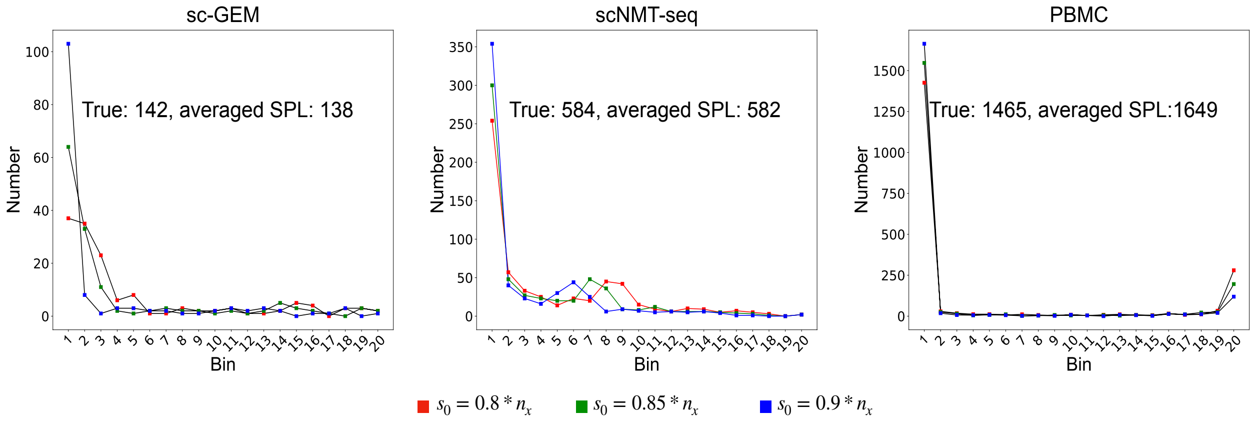


**Supplementary Figure 14.** Scree-Plot-Like (SPL) accurately estimated the shared cell number $s$ on scGEM, scNMT and PBMC data sets. We applied SPL by choosing $s_0 \in \{0.8, 0.85, 0.9\} * n_x$, repsectively, and used the averaged SPL results of three estimations as the final shared cell number $\hat{s}$.
